# Telomerase reverse transcriptase (TERT)-expressing cells mark a novel stem cell population in the adult mouse brain

**DOI:** 10.1101/2023.02.09.527879

**Authors:** Gabriel S. Jensen, Ashleigh N. Beaulieu, Caroline D. Curtis, Joshua Passarelli, Magdalena Blaszkiewicz, Seth Thomas, Trevor Morin, Jake W. Willows, Callie W. Greco, Ciara J. Brennan, Christopher Aniapam, Lydia Caron, Michele J. Alves, Matthew D. Lynes, Diana L. Carlone, David T. Breault, Kristy L. Townsend

## Abstract

Telomerase reverse transcriptase (TERT) is expressed by quiescent adult stem cells (ASC) in numerous adult murine and human tissues, but has never been explored in the adult brain. Here, we demonstrate that TERT+ cells in the adult mouse brain represent a novel population of multipotent ASCs that are localized to numerous classical neuro/gliogenic niches (including the ventricular-subventricular zone, hypothalamus, and olfactory bulb), as well as more recently described regions of adult brain plasticity such as the meninges and choroid plexus. Using a direct-reporter mouse line, we found that TERT+ cells expressed known neural stem cell markers such as Nestin and Sox2, but not markers of committed stem/progenitor cells, nor markers of mature neuronal or glial cells. TERT+ ASCs rarely expressed the proliferation marker Ki67, and *in vitro* TERT+ cells lost TERT expression when activated by growth factors, together indicating a quiescent phenotype similar to what has been observed in other tissues. When cultured, TERT+ cells behaved like neural stem cells by forming neurospheres, which could proliferate and become more metabolically active once stimulated by growth factors. TERT+ cells were observed in numerous brain niches, particularly near the ventricles and cerebrospinal fluid barriers, but notably, TERT+ cells were never observed in the hippocampus. Lineage tracing of TERT+ cells in adult transgenic mice (mTERTrtTA::oTET-Cre::RosamTmG) revealed large-scale expansion of TERT+ progeny and differentiation to diverse cell types in multiple brain regions. For example, lineage-traced cells expressed markers of mature neurons, oligodendrocytes, astrocytes, ependymal cells, and choroid epithelial cells, thus demonstrating the striking multipotency of this stem cell population in basal tissue turnover of the adult brain. Together, these data demonstrate that TERT+ cells represent a novel population of multipotent stem cells that contribute to basal plasticity and regeneration in the adult mouse brain.

**Graphical Abstract:** 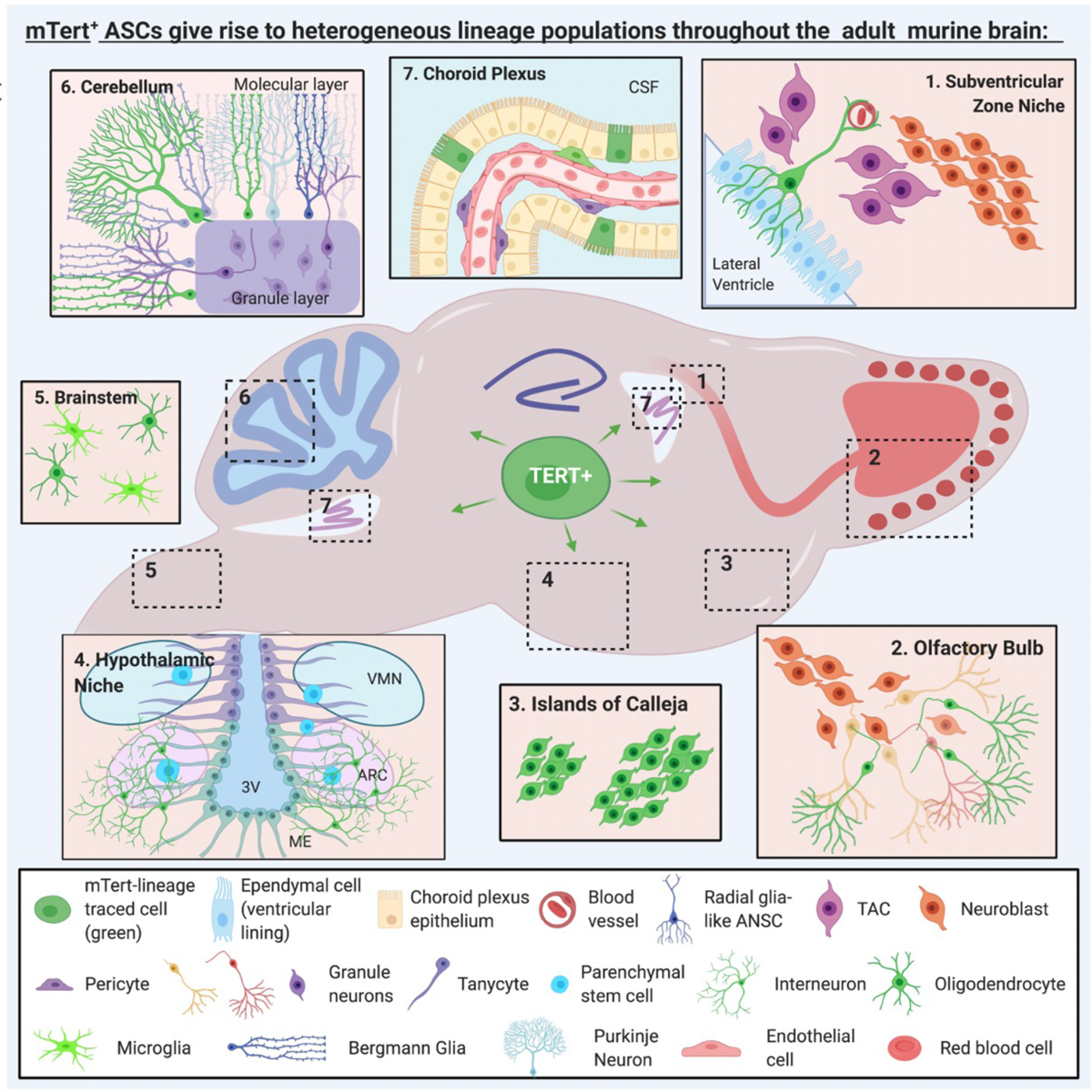

## Introduction

The adult mammalian brain retains cellular plasticity and the ability to regenerate certain cell types, but adult neurogenesis is thought to occur only in restricted patterns and discrete niches of the adult brain.^1^ Adult neurogenesis in mice is well-studied in the ventricular-subventricular zone (V-SVZ) of the lateral ventricles and the subgranular zone (SGZ) of the dentate gyrus (DG) in the hippocampus. However, numerous other adult brain anatomical niches have also demonstrated plasticity and capacity for neurogenesis, such as the hypothalamus, striatum, and amygdala.^2^ Adult neuro- and gliogenesis is supported by tissue stem cells, but a single stem cell population serving multiple brain niches has not yet been described.

Telomerase reverse transcriptase (TERT), the catalytic subunit of the enzyme telomerase, has previously been identified as a marker for slowly cycling or quiescent intestinal stem cells, as well as adult tissue stem/progenitor cells found in the testes, bone marrow, heart, thymus, spleen, liver, kidney, endometrium, heart, bone and adipose.^3–12^ However, until now TERT had never been investigated as a marker of adult stem cells (ASCs) in the brain. In both mice and humans, telomerase activity is high in the brain during development, but is dramatically reduced in the postnatal brain.^13, 14^ In mice, telomerase activity declines in the brain before adulthood, except in reportedly neurogenic regions such as the V-SVZ and olfactory bulb (OB).^9, 15, 16^ Numerous studies have pointed to TERT as a putative stem cell marker in the adult brain, but have stopped short of any confirmatory studies. For example, prior work revealed that telomerase activity was localized to cellular fractions containing Type A, B, and C neural progenitor cells (NPCs) from the adult mouse V-SVZ,^15^ and differentiation of embryonic stem cells into astrocytes or neurons induced loss of telomerase activity.^17, 18^

As mice age, they exhibit decreased TERT expression in the V-SVZ, which is accompanied by reduced rates of proliferation and neurogenesis, consistent with an aging-induced decline in the stem cell pool.^19^ Further evidence for the importance of telomerase activity in neurogenic niches is supported by decreased levels of V-SVZ neurogenesis in mice with telomerase RNA component (TERC) deletion. In these mice, stem cells in the V-SVZ are affected, while the proliferative ability of more committed/differentiated cells, such neuroblasts, was unaffected.^19^ Finally, in inducible TERT KO mice, tissue degeneration (including brain) and decreased V-SVZ neurogenesis were reversed following TERT reactivation.^20^ Together, these findings strongly suggested that TERT plays an essential role in adult stem cell maintenance and subsequently in adult neurogenesis, but we now provide new and compelling data to support these prior conjectures by describing TERT+ ASCs as multipotent stem cells in multiple anatomical niches of the adult mouse brain.

In the V-SVZ, adult neural stem cells (ANSCs) are important for replacing cells during basal tissue turnover and after injury, as V-SVZ-derived neuroblasts can reroute their typical migration to the OB to the site of cortical injury in order to aid in tissue repair.^21^ To better understand the future clinical potential of ANSCs to aid in repair or regeneration after brain injury, neurodegeneration, or normal aging requires a deeper mechanistic knowledge of neurogenesis and the potentially distinct populations of adult stem cells (ASCs) that regenerate various types of mature brain cells in different anatomical niches. In the absence of a specific and unique marker for ASCs in the brain, explorations have relied on a combination of cellular markers. Sex determining region Y-box 2 (Sox2), glutamate aspartate transporter 1 (GLAST), and glial fibrillary acidic protein (GFAP) are often used to identify ANSCs in V-SVZ and SGZ. Further complicating these analyses, each of these markers is also expressed by other cell types throughout the brain, reducing their specificity. (For example, GFAP and GLAST are expressed by mature glial cells in addition to stem cells.) We now provide new data to propose that TERT is a specific and novel marker for a population of multipotent ASCs in numerous niches of the adult mouse brain.

Here we demonstrate for the first time that TERT+ cells are observed in the adult mouse V-SVZ, hypothalamus, cerebellum, choroid plexus, and other regions. These TERT+ cells represent a rare, largely quiescent stem cell population that expresses several classical stem cell markers. We demonstrate that these cells possess the multipotent capability to proliferate upon activation and to regenerate multiple brain regions by differentiating into numerous mature cell types such as Purkinje neurons, olfactory sensory neurons, astrocytes, microglia, ependymal cells, and choroid plexus epithelial cells.

## Materials and Methods

### Mice

All procedures were approved by the University of Maine and The Ohio State University IACUCs. Mice were maintained on a 12-hr light/dark cycle, and food and water were provided *ad libitum*. Tg(Tert-GFP)22Brlt (*mTERT*-GFP) mice were described previously.^3^ For lineage tracing studies, *mTert*-rtTA::otet-Cre::R26R^flox(mTmG)^ mice were generated by crossing *mTERT-*rtTA mice^4^ to otet-Cre::R26R^flox(mTmG)^ mice, which were generated by crossing B6.Cg-Tg(tetO-cre)1Jaw/J mice (The Jackson Lab) to B6.129(Cg)-Gt(ROSA)26Sor^tm4(ACTB-tdTomato,-EGFP)Luo^ mice (The Jackson Lab). Recombination was induced via 2 mg/ml doxycycline (Sigma Aldrich) in 50 mg/ml sucrose water over the course of 2 days to several weeks, as indicated for the ‘pulse’ period to mark TERT+ cells green. To ‘chase’ this signal in lineage tracing experiments, the doxycycline water was removed, and normal water was provided for various chase periods of days to weeks. Both male and female animals were utilized for most experiments (details in each Figure/Results section).

### Brain Collection and Processing

Mice were anesthetized via intramuscular injection of 320 mg/kg body weight Ketamine and 24 mg/kg body weight Xylazine in 0.9% saline and then euthanized via transcardial perfusion with 1X PBS (phosphate buffered saline) followed by Histochoice Fixative or 4% paraformaldehyde (PFA). Brains were dissected out and post-fixed overnight at 4°C, placed in 15% sucrose and 30% sucrose at 4°C for 2 days each to cryopreserve, and embedded in OCT. Brains were sliced at 7-10 µm on a CM1950 or CM3050 S cryostat (Leica) at −20°C and serial sections were placed into each Superfrost slide and stored at −20C for downstream immunostaining.

### Immunostaining

Prior to the immunostaining, all the slides were brought to room temperature and post-fixed with ice-cold acetone for 15min. Slides were washed for 5min in 1X IHC Select TBS Rinse Buffer (Millipore), shaken at 60rpm at room temperature (RT) between each step. Permeabilization was performed with either 0.3% Triton X-100 (nuclear antigens) or 0.3% Tween-20 (cytoplasmic antigens) for 10min at RT. Antigen retrieval was performed with 1X DAKO Antigen Retrieval Solution in the microwave (on low power) for 20 minutes. Slides were incubated in 0.3% Typogen Black in 70% EtOH for 20min at RT (to quench autofluorescence), blocked for 20 min at 37°C with Millipore Blocking Reagent, and incubated with primary antibody overnight at 4°C. Antibody concentrations are listed in the Key Resource Table. The Alexa Fluor secondary antibodies were incubated for 10m at RT, then counter-stained with DAPI (100ng/mL) for 5min to visualize cellular nuclei. Epifluorescence images were captures on Nikon E400 microscope with a Hamamatsu Flash 2.0. Confocal images were captures on either SP8 (Leica), Stellaris 5 (Leica), or LSM 900 (Zeiss) for co-expression analysis.

### Fluorescence Activated Cell Sorting (FACS)

To obtain single cell suspensions of adult mouse brains, mice were euthanized via CO_2_ asphyxiation followed by cervical dislocation. Brains were dissected out and washed with artificial cerebrospinal fluid (ACSF; Ecocyte). After the tissue was cut into smaller pieces with a razor blade in a petri dish containing ice-cold ACSF with 1mg/mL pronase (Millipore-Sigma), it was incubated for 60-75min at 37°C and 90rpm in a shaking water bath incubator. Every 5-10 minutes tubes were vortexed and then returned to shaking incubator. Samples were centrifuged for 4min at 1600rpm and supernatant decanted. Samples were resuspended with ACSF with 5% fetal bovine serum (FBS) and incubated for 15min at 37°C and 90 rpm. Trituration of each sample was performed with glass Pasteur pipettes with approximately 600μm, 300μm, and 150μm openings, followed by centrifugation for 10min at 300 x g and discard of supernatant fraction. Debris removal solution (Miltenyi) was utilized to remove debris and myelin from the samples. For separation of CD45 cells, the cell suspension was treated with 1X PBS with 0.5% BSA and 2% FBS for 20 min at 4°C, centrifuged for 5min at 1800rpm and resuspended in 1X PBS with 5% FBS and 5mM EDTA containing antibodies for 20min at 4°C. The cell suspension was washed twice with 1X PBS with 0.5% BSA and 2% FBS and resuspended in ACSF with 10% FBS. Cells were first sorted for CD45 signal, and negative cell population were then sorted for endogenous membrane GFP fluorescent signal via BD FACSAria II (Jackson Laboratory, Bar Harbor, ME) or BD FACSAria III (OSUMC, Columbus, OH). GFP+/CD45-cells were sorted into Trizol (Ambion) and frozen at −80°C for qRT-PCR or Neurosphere Basal Medium supplemented with Proliferation Supplement (STEMCELL Technologies) as described below. Additionally, PSA-NCAM Antibody, anti-human/mouse/rat, APC (Milteny, Ref. 130-120-437) was utilized to sort neurons through BD FACSAria III for further qPCR downstream application.

### Magnetic Activated Cell Separation (MACS)

*mTERT-*rtTA mice^4^ to otet-Cre::R26R^flox(mTmG)^ mice received doxocycline as described above for 5 days and were euthanized at day 5 via CO_2_ asphyxiation followed by cervical dislocation. Dissociation of the brain was carried out with the Adult Brain Dissociation Kit for mouse and rat (Milteny Biotec, Ref.130-107-677) using the gentleMACS™ Octo Dissociator with Heaters, as per the manufacture’s recommendations. Briefly, brains were dissected out, washed with cold D-PBS, added to the enzyme mix 1, and diced to smaller pieces. The enzyme mix 2 was added and samples were incubated with gentleMACS Program 37C_ABDK_01. Samples were applied through a cell strainer (70µm) and 10ml of DPBS was added during the process. After samples being centrifuged 300×g for 10 minutes at 4°C, the debris removal step was performed, followed by the red blood cells removal. Next, the cells were resuspended in cold DPBS pH 7.2, 0.5% BSA and 2 mM EDTA, added to 10ul of CD45 MicroBeads (Milteny, Ref. 130-052-301) per 10^7^ total cells. After 15min of incubation at 4-8°C, the cell suspension was washed with buffer and proceeded to the magnetic separation. Cells suspension was applied to MACS LS Column and MACS Separator and total effluent was collected as CD45 negative population. The CD45-cells were washed and resuspended in DPBS pH 7.2, 0.5% BSA and 2 mM EDTA added to Anti-GFP antibody (1:500, Abcam, Ref. ab6556) for 15min at 4-8°C. The magnetic labeling using Anti-Rabbit IgG MicroBeads (Milteny, Ref.130-048-602) was carried out to select GFP+ cells following the magnetic separation with LS columns. Cell suspension was resuspended either in Neurosphere Basal Medium containing Proliferation Supplement (STEMCELL Technologies) 20ng/mL hrEGF (STEMCELL Technologies), 10ng/mL hrbFGF (STEMCELL Technologies), and 0.002% Heparin (w/v) (STEMCELL Technologies), or in RNeasy Micro Kit (Qiagen, Ref. 74004) lysis buffer for qPCR applications.

### Neurosphere Formation Assay

*In vitro* neurosphere culturing experiments were performed according to methods from ^22^. Briefly, after FACS or MACS sorting, cells were cultured into 96-well low-attachment plates containing NeuroCult Basal Media (Mouse and Rat; STEMCELL Technologies) with Proliferation Supplement (STEMCELL Technologies), 20ng/mL hrEGF (STEMCELL Technologies), 10ng/mL hrbFGF (STEMCELL Technologies), and 0.002% Heparin (w/v) (STEMCELL Technologies). Both GFP+ and GFP-cell population were grown at 37°C with 5% CO_2_ and 90% relative humidity for 14 days or longer. Media was changed every 5-7 days. For certain experiments, cells were imaged every three days for a total of 22 days.

### EdU Labeling

Neurospheres were removed from 96-well plates and pipetted into 8-well Seahorse XF HS miniplates (Agilent) coated with 15 µg/mL poly-L-ornithine and 10µg/mL laminin. Plates were spun at 200 x g for 2 minutes for adherence. Cells were incubated in 5µM EdU (ThermoFisher Scientific) in NeuroCult proliferation media for 24hr. Cells were fixed with 4% PFA (pH 7.4) for 15min, washed twice with 3% BSA in 1X PBS, and permeabilized with 0.5% Triton X-100 in 1x PBS for 20min at RT. Cells were washed once more and incubated in Click-iT EdU reaction buffer (ThermoFisher Scientific). Cells were then washed once with PBS and imaged.

### RNA Extraction and qRT-PCR

Total RNA extraction was performed using either Zymo Direct-zol RNA Microprep kit (Zymo) or RNeasy Micro Kit (Qiagen, Ref 74004) for GFP+ cells population after MACS. RNA quality and concentration were assessed using a Nanodrop One (ThermoFisher). Reverse transcription was performed with the High-Capacity cDNA Reverse Transcription Kit (ThermoFisher) according to the manufacturer’s instructions. qRT-PCR reactions were carried out using Universal SSA SYBR Green (ThermoFisher) on a CFX384 qPCR System (Bio-Rad). The final volume for each reaction was 10μl with 100nM of corresponding gene specific primers, and 1μl of total cDNA. No-template controls were run alongside each primer pair. All reactions were carried out in duplicates. The thermal cycling was initiated at 95°C for 20s followed by 40 cycles of 5s at 95°C and 30s at the optimal annealing temperature for each gene. Dissociation curves were analyzed at the end of each run for product verification. The mean Ct-values for each sample were normalized using cyclophilin as a housekeeping gene. Primer sequences are found in Key Resources Table 1.

### Seahorse Assays

Following the MACS separation and sorting of GFP+ cells, these were cultured into either 96-well low-attachment plates containing Neurosphere Basal Medium containing Proliferation Supplement (STEMCELL Technologies) 20ng/mL hrEGF (STEMCELL Technologies), 10ng/mL hrbFGF (STEMCELL Technologies), and 0.002% Heparin (w/v) (STEMCELL Technologies), or directly into 8-well Seahorse XF HS miniplates (Agilent) coated with 15 µg/mL poly-L-ornithine and 10µg/mL laminin. They were kept at 37°C with 5% CO_2_ and 90% relative humidity and media was added every other day. After XX days, neurospheres were dissociated with Accutase and 2.5×10^3^ were seeded in each well of 8-well Seahorse XF HS miniplates coated as described above. The media was replaced for the XF DMEM ph 7.4 with 1% (v/v) of XF Glucose, XF pyruvate, XF L-Glutamine and cells were prepared for the Mito Stress Assay (Agilent) as per manufacture’s recommendations. The final concentration of the compounds used was 1.5µM of Oligomycin, 1.5 µM of FCCP, 0.5 µM of Rot/AA. The assay was performed Agilent Seahorse XF HS Mini Analyzer.

### Live Imaging (IVIS-CT)

For acquisition of bioluminescent images, mice were sedated with 2% isoflurane. D-Luciferin (Perkin Elmer) was diluted to 3 mg/100 µL in normal saline and 0.6 mg of D-Luciferin was administrated intraperitoneally. 15 minutes after injection, an IVIS-Spectrum CT imaging system equipped with a CCD camera (Caliper Life Sciences) was used for 3D bioluminescence using the standard single-mouse resolution MicroCT and bioluminescence was projected onto this 3-dimensional data using Living Image software (Caliper Life Sciences). Mice were then sacrificed and dissected and imaged again approximately 20 minutes after the D-Luciferin injection.

### Intracerebroventricular (i.c.v.) Administration of Mitotic Inhibitor Ara-C

13 weeks old mice were administered with 2mg/ml doxycycline in the drinking water for a 4-week pulse to activate mTERT expressing cells with GFP. Prior to doxycycline removal (06 days), they received cytosine-β-D-arabinoside (2%; Ara-C) in 0.9% saline or 0.9% saline alone into the right lateral ventricle via a mini-osmotic pump (ALZET) with cannulas implanted. Brain cannulation occurred at the following coordinates: anterior/posterior (A/P), − 0.5mm; medial/lateral (M/L), −1.1mm; dorsal/ventral (D/V), −2.5mm (relative to Bregma and the surface of the brain; M/L coordinates are directed laterally to the right). After 6 days of Ara-C infusion, the mice were perfused, and the brains were collected as described above.

### Optical Clearing and Immunostaining/Imaging of Brain Sections

Adult male mTert-rtTA(+/+)::oTet-Cre(+/-)::Rosa-mTmG(+/+) mice (N=3) and Cre-, negative control, mice (mTert-rtTA(+/+)::oTet-Cre(-/-)::Rosa-mTmG(+/+), N=1) were transcardially perfused with 1X PBS and 4% PFA after a 3-week, 11-day doxycycline pulse-chase. Brains were dissect out and post-fixed in 4% PFA overnight at 4°C followed by 1hr at RT. Brains were washed twice in 1X PBS for 60min, cut into 500 μm sagittal sections using brain matrices, transferred to microcentrifuge tubes containing 1X PBS and pre-treated with methanol as described by Renier *et al.*^23^ Next, samples were incubated in 1X PBS/0.2% TritonX-100/20% DMSO/0.3M glycine overnight at 37°C. The blocking was performed by incubating these samples in 1X PBS/0.2% TritonX-100/10% DMSO/6% donkey serum at 37°C for 3 days on an orbital shaker. Wash steps were performed twice in buffer containing 1X PBS/0.2% Tween-20 with 10µg/ml heparin for 1h at 37°C. Rabbit anti-GFP Alexa Fluor 488 (1:500) was added into 1X PBS/0.2% Tween-20/10µg/ml heparin/5% DMSO/3% donkey serum and incubated at 37°C for 2 days. Samples were then washed 3 times for 1 hour at 37°C on orbital shaker, and then once a day for 2 days. Tissues were then cleared with the iDISCO method.^23^ Cleared brains were refractive index matched with, and imaged while submerged in, dibenzyl ether which is corrosive to most objective lenses and most plastics. To mitigate this, we took a glass bottom dish and coated any interior plastic with quick drying silicone elastomer (Kwik-Sil^TM^). Cleared brains were imaged on a Leica Stellaris 5 confocal microscope with a white light laser (ex. 495, em. 505-600) intensity set to 20%. Images were captured with a 10X dry objective at 1024×1024 resolutions with pinhole set to 1 AU. Line scan speed was set to 600 Hz and lines were averaged 3 times per frame. Photons were captured with Power HyD S detectors; gain set to 20%. Images were captured as Z-μm step-size (84 images per stack) Z-stacks were tiled together to form a complete image of each brain slice. These were then Z-maximum intensity projected into a comprehensive 2D image.

### Statistical Analyses

All plots represent mean ± SEM. Statistical calculations were carried out in Excel or GraphPad Prism 9.0, utilizing the ANOVA or Student’s T-test as indications of significance. For all figures, \**P*<0.05, \*\**P*<0.01, \*\*\**P*<0.005, \*\*\*\**P*<0.001.

### Data availability

The authors confirm that the data supporting the findings of this study are available within the article [and/or] its supplementary material. Raw data were generated at University of Maine and The Ohio State University. Derived data supporting the findings of this study are available from the corresponding author on request.

## Results

### TERT+ cells in the brain of adult mTert-GFP direct reporter mice are localized to the V-SVZ and other brain regions with reported adult neural plasticity

To characterize the location and quantity of mTERT-GFP+ cells in the adult mouse brain, we first examined a neurogenic-enriched area including V-SVZ, SGZ, and hypothalamus^3^ from *mTert*-GFP direct reporter mice as illustrated in Figure 1A. Our analysis revealed localization of TERT-GFP+ cells to most of the classical neurogenic niches, as well as newly identified plastic niches such as the choroid plexus (ChP) and meninges^24, 25^ (Figure 1B). TERT+ cells were detected at low numbers, as expected for quiescent ASCs given their function as a reservoir that can proliferate upon activation, and as has been seen previously in other tissues like the gut.^3,10^ Most of the observed TERT+ cells were identified in the V-SVZ, LV ChP, D3V, hypothalamus, and meninges, and there were significantly more TERT+ cells in the meninges when compared to the V-SVZ (Figure 1B, S1A). TERT+ cells were observed less frequently in areas such as cerebral cortex (Figure 1B). Interestingly, no TERT+ cells were ever identified in the DG of the hippocampus across any of the 20+ cohorts of mice analyzed for this manuscript (representative data in Figure 4). Across several Bregma coordinates (9 in total) of the neurogenic region-enriched areas, our analysis showed that TERT+ cells were found at similar numbers with no significant differences in TERT+ cells per niche, within each Bregma region (Figure 1C-D). Fluorescence activated cell sorting (FACS) of TERT-GFP+ cells from entire *mTert*-GFP brains revealed that only 0.1-0.3% of brain cells were TERT+ (Figure S1A), fitting with the frequency of TERT+ ASCs found in other tissues like gut^3,10^. Taken together, TERT+ cells are found in low numbers throughout various niches of the adult mouse brain that are known to exhibit adult neural plasticity and/or neurogenesis.

**Figure 1.**
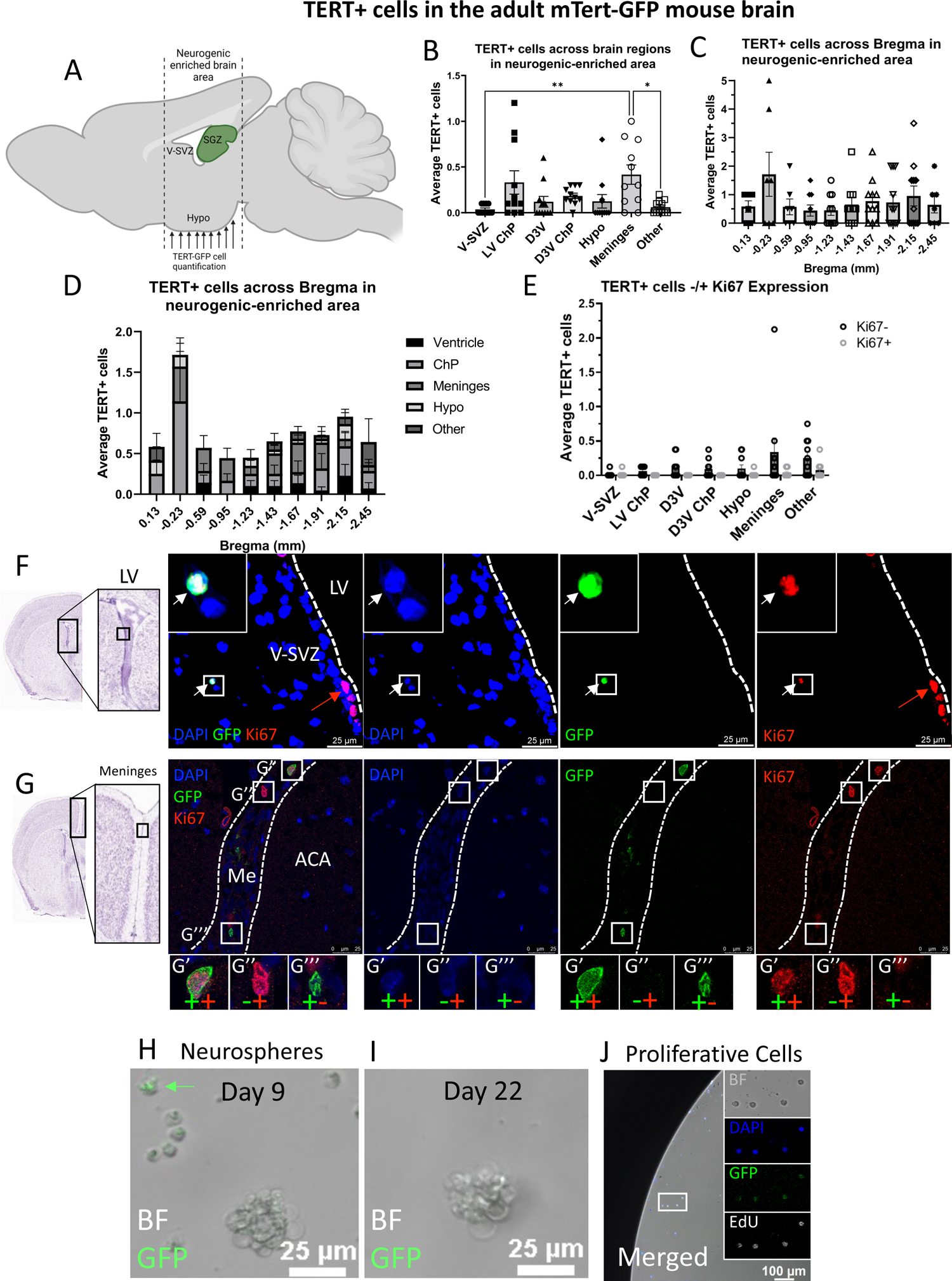
TERT+ cells in the adult mTert-GFP mouse brain are quiescent stem cells localized to both well-studied and novel regions of adult brain plasticity. (A) Mouse brain illustration of the neurogenic-enriched brain area used for immunostaining with arrows indicating coronal planes used for TERT-GFP+ cell analysis. (B) Quantification of TERT + cells per region identified via immunostaining in adult mTert-GFP mouse brains. Cell quantification is graphed as the average number of TERT+ cells per niche, per mouse. 8-20 coronal brain sections per mouse were analyzed from *N* = 11 mice (*N* = 5 males and *N* = 6 females) at 12-weeks of age. Dots represent individual animals (C) Quantification of TERT+ cell numbers across Bregma coordinates through the neurogenic enriched brain area. Cell quantification is graphed as the average TERT+ cell number across 2 coronal brain sections per depth, per mouse. *N* = 6-11 mice per depth. Dots represent individual animals. (D) Quantification of TERT+ cells per region across Bregma coordinates through the neurogenic enriched brain area. Cell quantification is graphed as the average number of cells per brain region across *n =* 2 brain sections for each of *N =* 6-11 mice. (E) Quantification of co-staining for TERT-GFP and Ki67 in mTert-GFP mouse brains. Cell quantification is graphed as the average number of cells per brain region across *n* = 8 brain sections for each of *N* = 5 males and 5 females at 12-weeks of age. (F) Co-immunostaining of TERT+ cell with the proliferation marker Ki67 in the V-SVZ. White arrow indicates TERT+Ki67+ cell. Red arrow indicates TERT-Ki67+ cells. 8 brain sections were analyzed for each of *N* = 10 mice (*N* = 5 males and 5 females) at 12-weeks of age. (G) Representative image of co-immunostaining with Ki67 in the meninges of mTert-GFP mice. Green or red “+” or “-” indicates that the cells in indicated area were GFP+/GFP-, or Ki67+/Ki67-, respectively. Dots indicate individual animals. 8 sections per brain were analyzed from *N* = 8 mice (*N* = 4 males and *N* = 4 females) at 12-weeks of age. (H-I) Representative image of neurosphere formation of GFP+CD45-cells with fluorescence analyzed on Day 9 (H) and Day 22 (I). *N* = 6 animals at 12-20-weeks of age. (J) EdU incorporation in TERT+ cells cultured with proliferation media containing EdU for 24 hours starting at day 9 (*N* = 3 of 4 mice, *n* = 1 well per brain). Scale bars are 100µm. Insets show 4x digital magnification of indicated area. Bar graphs show mean cells per brain section, per animal ± SEM with individual data points per animal. Grouped bar graphs show mean cells per mouse, per brain section, at each depth ± SEM. Significance was determined via one-way ANOVA (\**P* < 0.05, \*\**P* < 0.01). V-SVZ: ventricular-subventricular zone, LV: lateral ventricle, ChP: choroid plexus, D3V: dorsal 3rd ventricle, Hypo: hypothalamus, Me: meninges, ACA: anterior cingulate area, BF: brightfield, EdU: 5-ethynyl-2’-deoxyuridine.

### TERT+ cells in the adult mTert-GFP mouse brain are mostly quiescent and not proliferative

Quiescent populations of stem cells are expected to slowly divide or proliferate, to prevent exhaustion of the stem cell pool. Previously, it had been reported that TERT+ adult tissue stem cells in the intestines are slowly cycling and rarely express the proliferation marker Ki67.^3,10^ Expression of TERT may act to restore telomere length after cell division and prevent senescence into adulthood, which is a defining characteristic of stem cells. To assess the proliferative capacity of TERT+ cells in the adult mouse brain, we co-immunostained with the proliferation marker Ki67 in the neurogenic-enriched area (the region shown in Figure 1A) of the adult brain in *mTert*-GFP mice.^3^ Figure 1E shows that TERT+Ki67+ double positive cells were mostly identified in the V-SVZ, hypothalamus, and meninges, with fewer in other brain regions. TERT+Ki67+ cells were not found in the LV ChP, D3V, or D3V ChP (Figure 1E).

Of all TERT+ cells analyzed across all brain sections, only 14.9% of TERT+ cells co-expressed Ki67 (Figure 1E, S1B). In addition, we observed every iteration of co-stained cells: TERT+Ki67+ (Figures 1F, 1G’), TERT+Ki67-(Figure 1G’’), and TERT-Ki67+ (Figure 1G’’’), indicating that TERT expression is not cell-cycle specific, yet is *<u>rarely</u>* expressed by actively dividing cells. Our results demonstrate that TERT+ cells in the adult brain possess quiescent stem cells features, as only a small percentage of them express the proliferative marker Ki67.

Previous studies have identified subpopulations of TERT+ cells as CD45+ immune cells originating from bone marrow in various murine tissues.^5, 10^ To determine whether TERT+ cells in the adult murine brain also included a population of CD45+ immune cells that may have migrated to the brain from the periphery, we quantified both TERT+CD45+ and TERT+CD45-cells throughout the adult mouse brain by immunostaining (Figures S1C-S1E) and by FACS cell sorting (Figures S1F-I). These data revealed that 53.6% of TERT+ sorted cells expressed CD45 (Figure S1F). TERT+CD45+ cells were identified across several brain regions, especially in the meninges, V-SVZ, D3V and D3V ChP. Interestingly, no TERT+CD45+ cells were identified in the hypothalamus by immunostaining (Figure S1E).

FACS analysis of *mTert*-GFP brains, in which CD45+ cells were gated separately to exclude them, revealed that 99% of brain cells were CD45- and that of these non-immune cells, only 0.1% were TERT-GFP+ (Figures SG-I). It is important to note that TERT+/CD45-cells may be a subpopulation of resting microglia expressing low levels of CD45 ^26^ which may have maintained TERT expression into maturity.

### TERT+ cells form neurospheres in culture

The ability of neural stem cells to form neurospheres in culture is a standard assay for characterization of brain stem cells.^27^ To analyze the neurosphere-forming capabilities of a pure population of TERT+ stem cells from the adult mouse brain, TERT+CD45-stem cells were sorted by FACS (to eliminate TERT+ immune cells) from GFP direct reporter mice and plated into neurosphere growth media containing epidermal growth factor (EGF) and basic fibroblast growth factor (bFGF), as described previously.^22^ The strategy for isolation of GFP+CD45-cells is outlined in Figure S1G-I. We found that isolated TERT+ cells did form classical neurospheres in culture (Figure 1H). GFP-cells sorted from *mTert-*GFP animals were used as negative controls (Figure S1J). Individual cells retained GFP expression in culture, while mature neurospheres exhibited a loss of GFP expression in culture over time (Figures 1H-J), likely due to their activation state due and loss of quiescence by the presence of growth factors in the culture media.

In addition, TERT+ neurospheres incorporated 5-ethynyl-2’-deoxyuridine (EdU) after 24 hours in culture, indicating a potential for proliferation during the activation period in the presence of growth factors (Figures 1J, S1K). Since a subset of the FACS-sorted cells lost GFP expression *in vitro*, we quantified EdU+/GFP+ cells and observed that 49.5% ± 5.6 of EdU+ cells also co-expressed GFP+ (Figure S1L). Together these results demonstrate that TERT+ cells in the adult mouse brain are a population of rare, neurosphere-forming stem cells with the potential for proliferation *in vitro* and that GFP expression is transiently maintained during differentiation but is turned off after activation and neurosphere formation, likely due to loss of TERT expression in that cell state.

### TERT+ cells express markers of quiescent stem cells, but not of activated stem cells or committed neuronal progenitors

There is currently no single, specific marker for qASCs in the adult brain. This is due to the fact that all previously identified stem cell markers are also expressed by various non-stem cell types.^28^ For this reason, we investigated the expression profile of the classical neural stem cell markers Sox2, Nestin, and EGFR in TERT+ cells from adult *mTert*-GFP mouse brains. For reference, the neuronal lineage markers are shown in Figure 2A. We found populations of TERT+Sox2+ and TERT+Sox2-cells throughout all major TERT+ niches except for the V-SVZ and D3V, which housed no TERT+Sox2+ double positive cells (Figures 2B-D). Analysis of the total number of TERT+ cells across all brain sections revealed that 33.3% of TERT+ cells expressed Sox2 (Figure S2A). Interestingly, we only uncovered a single TERT+Nestin+ cell (Figure 2E, S2B) compared to other 9 TERT+Nestin-cells in the same brain region, indicating that only 10% of TERT+ cells were Nestin+ (Figure 2E-F, S2B) across all the cohorts analyzed for this study. These results show that only a subset of TERT+ cells expressed the markers Sox2 or Nestin and therefore may represent a novel brain stem cell pool, potentially with some overlap with the Sox2+ stem cells that have been investigated previously.

**Figure 2.**
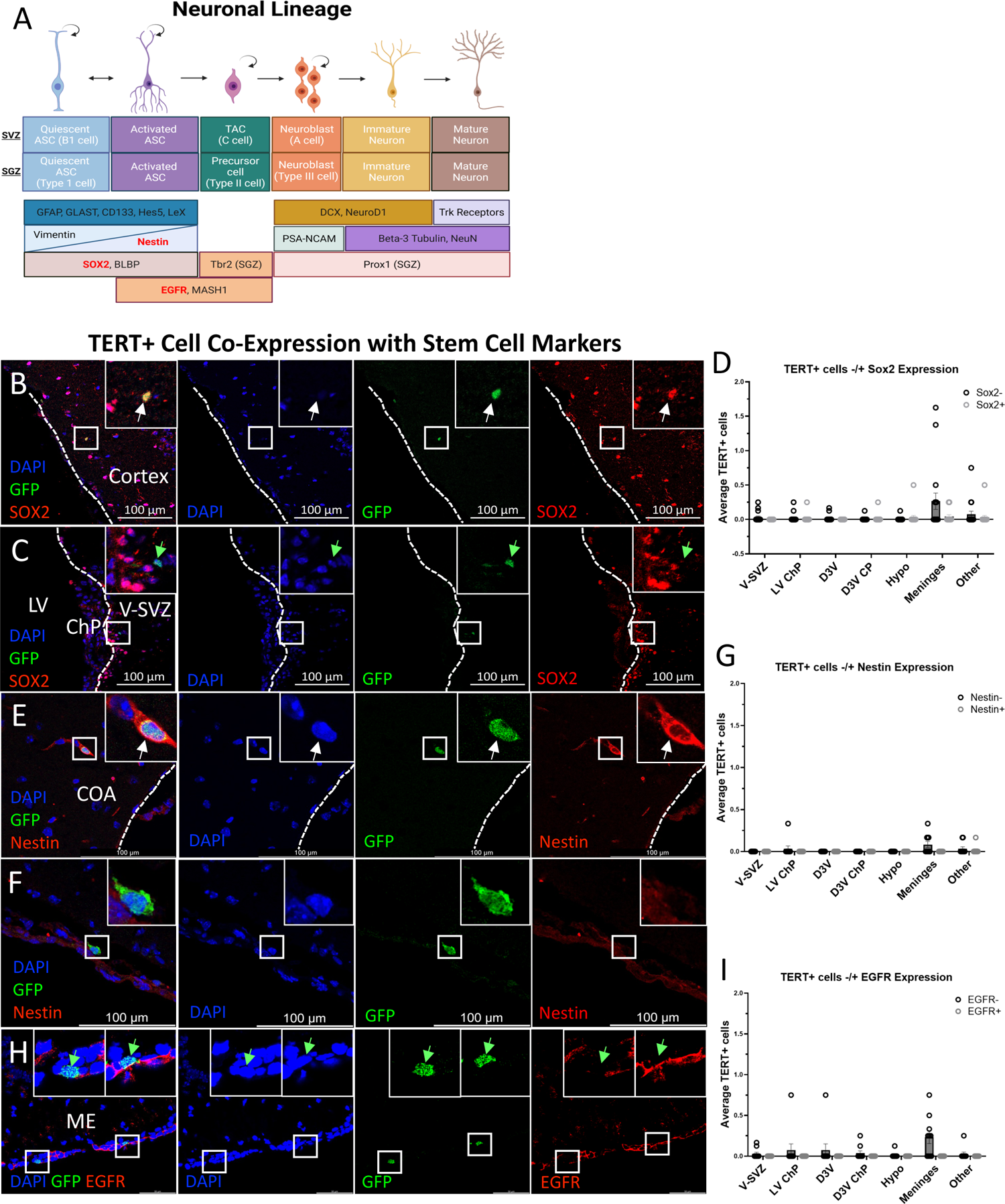
TERT+ cells express markers of quiescent stem cells, but not of activated stem cells, neuronal precursors, or mature neurons. **(A)** Schematic depicting neuronal lineage markers from qASCs to mature neurons in the adult mouse brain. Markers in red text were stained for in this figure. SGZ-specific markers are denoted (SGZ). **(B-C)** Representative images of Sox2 staining in the adult mTert-GFP mouse brain. TERT+Sox2+ cells were identified in *N* = 7 of 17 mice (*N* = 4 of 8 males and 3 of 9 females), *n* = 4-8 brain sections per mouse (B). TERT+Sox2-cells were identified in *N* = 10 of 17 mice (*N* = 4 of 8 males and 6 of 9 females), *n* = 4-8 brain sections per mouse (C). **D)** Quantification of immunofluorescent co-staining analysis of the stem cell marker Sox2 in adult mTert-GFP mice. Cell quantification is graphed as the average number of cells per brain region across *n* = 4-8 brain sections per mouse from *N* = 17 mice (*N* = 8 males and *N* = 9 females) at 12 weeks of age. **(E-F)** Representative images of Nestin staining in the adult mTert-GFP mouse brain. The rare TERT+Nestin+ cell was identified in *N* = 1 of 10 mice (*N* = 1 of 4 males and 0 of 6 females), *n* = 4-6 brain sections per mouse (E). TERT+Nestin-cells were identified in *N* = 5 of 10 mice (*N* = 3 of 4 males and 2 of 6 females), *n* = 4-8 brain sections per mouse (G). **(G)** Quantification of immunofluorescent co-staining analysis of the stem cell marker Nestin in adult mTert-GFP mice. Cell quantification is graphed as the average number of cells per brain region across *n* = 4-8 brain sections per mouse from *N* = 4 males and *N* = 6 females at 12 weeks of age. **(H)** Representative image of EGFR staining in the adult mTert-GFP mouse brain. TERT+EGFR-cells were identified in *N* = 8 of 10 mice (*N* = 3 of 4 males, *N* = 5 of 6 females), *n* = 4-6 brain sections per mouse. **(I)** Quantification of immunofluorescent co-staining analysis of the stem cell marker Nestin in adult mTert-GFP mice. Cell quantification is graphed as the average number of cells per brain region across n=4-6 brain sections per mouse from *N* = 4 males and *N* = 6 females at 12 weeks of age. Insets show indicated area at 3x digital zoom. White arrows indicate co-stained cells, green arrows indicate cells that express only TERT. Scale bars are 100µm except Figure H (50µm). Bar graphs show mean cells per mouse, per brain section, at each brain region ± SEM. All images taken on SP8 or LSM 900 confocal microscopes. V-SVZ: ventricular-subventricular zone, LV: lateral ventricle, ChP: choroid plexus, COA: cortical amygdalar layer, ME: median eminence.

To determine whether TERT+ cells could express markers of activated stem cells or differentiated cell types, we performed co-immunostaining. The activated adult stem cell marker EGFR, which expressed by active ASCs (aASCs), TACS (V-SVZ), and Type II cells (DG), was never expressed by TERT+ cells in any brain region (Figures 2H-I, S2C). Additionally, TERT+ cells never expressed the more differentiated neuroblast marker DCX or the mature neuron marker neuronal nuclear protein (NeuN) (Figures S2D-G). In addition, bone morphogenetic protein (BMP) receptor 1A (BMPR1A) which is a protein in the V-SVZ and SGZ that allows for stem cells to respond to neuro/gliogenic BMP signals^29^ was identified in only a single TERT+ cell (Figure S2H). Taken together, these results indicate that TERT+ cells express quiescent ASC markers but not markers of activated/committed progenitors or mature cell types.

### TERT+ cells do not express markers of glial progenitors or mature glia

W subsequently performed immunostaining for markers of glial precursor cells (GPCs) and mature glia, to determine if TERT+ cell types could be non-neuronal. GPCs expressing neuron-glial antigen 2 (NG2) are found throughout the adult brain and likely arise from Sox2+ and Nestin+ stem cells.^30^ Figure 3A depicts the changes in gene expression across glial cell differentiation, for reference. Co-immunostaining revealed no co-expression of TERT with any glial markers including NG2, the oligodendrocyte-lineage marker oligodendrocyte transcription factor 2 (OLIG2), nor the astrocyte-lineage marker GFAP (Figures 3B-H). Interestingly, although GFAP has been described as expressed on brain stem cells^31^, we did not see GFAP marking TERT+ ASCs, again indicating these may be a novel and distinct population of brain stem cells. Although, images using the OLIG2 antibody in the V-SVZ confirmed the labeling of OLIG2+ cells in other brain regions including the V-SVZ (Figure S3A), it was found not co-expressed with TERT+ cells. These data indicate that TERT+ cells in the murine brain are not glial-committed precursors, mature astrocytes, or mature oligodendrocytes.

**Figure 3.**
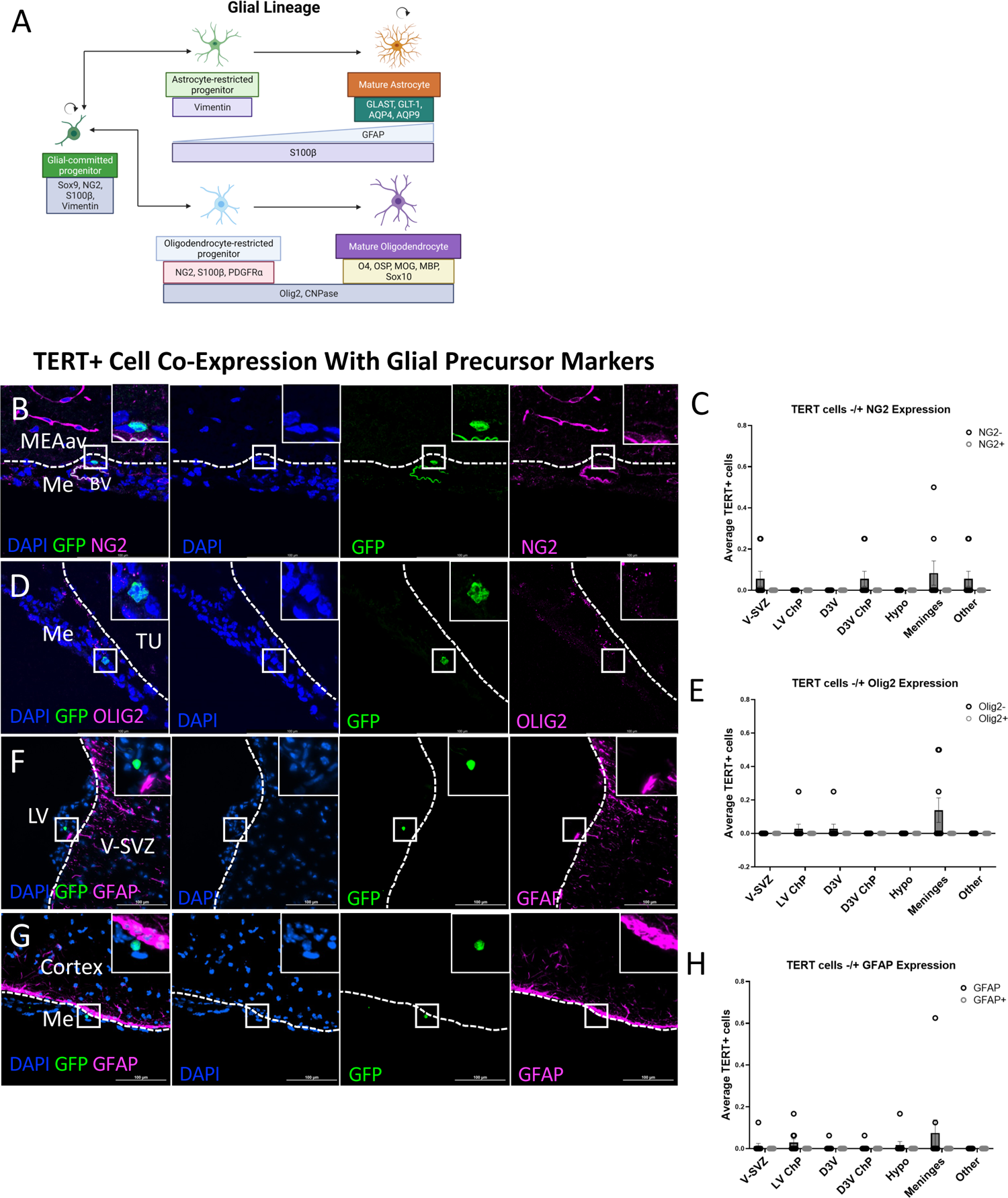
TERT+ cells do not express markers of glial-committed progenitors or mature glia. **(A)** Schematic depicting glial lineage markers from glial-committed progenitors to mature glial cell types in the adult mouse brain. Markers with red text were stained for in this figure. **(B-C)** Immunostaining in the brains of adult mTert-GFP mice for the glial precursor cell marker NG2 identified no TERT+NG2+ cells (*N* = 5 males, 5 females, *n* = 4 sections per brain analyzed). **(D-E)** Immunostaining in the brains of adult mTert-GFP mice for the progenitor/ mature oligodendrocyte marker OLIG2 identified no TERT+Olig2+ cells (*N* = 5 males, 5 females, *n* = 4 sections per brain analyzed). **(F-H)** Immunostaining in the brains of adult mTert-GFP mice for the marker of immature/ mature astrocytes as well as qANSCs identified no TERT+GFAP+ cells (*N* = 5 males, 5 females, *n* = 4 sections per brain analyzed). Insets show indicated area at 3x digital zoom. Scale bars are 100µm. MEAav: media amygdalar nucleus, anteroventral part, Me: meninges, TU: tuberal nucleus, LV: lateral ventricle, V-SVZ: ventricular-subventricular zone. Bar graphs show mean cells per mouse, per section at each brain region ± SEM.

### TERT+ stem cells lineage trace to multiple anatomical regions of the adult mouse brain

Based on the premise that TERT+ cells are ASCs, we tested the hypothesis that TERT+ cells would proliferate, differentiate, and migrate within the adult mouse brain using an inducible lineage tracing mouse system.^32^ Lineage tracing allows for an indelible tag, such as the bioluminescent luciferase or the fluorescent protein GFP, to be expressed by each TERT+ cell during a ‘pulse’ period with delivery of the tetracycline derivative doxycycline (‘dox’) in the drinking water. A ‘chase’ period after dox administration ends results in the discontinuation of TERT cell labeling but allows the tracing of these newly labeled cells as they migrate, proliferate, or differentiate into mature cell types. Lineage tracing was initially performed with *mTert*-rtTA::oTet-Cre::LSL-FLUC mice, which allowed whole body and whole brain small animal imaging of luciferase signals. In these mice, the 4.4kb portion of the *mTert* promoter drove expression of reverse tetracycline transactivator (rtTA), which activated Cre recombinase expression in the presence of dox. This allowed rtTA interaction with the tet-operator upstream of the Cre open reading frame. Cre mediated recombination of the bioluminescent (luciferase) lineage tracing alleles resulted in constitutive expression of the reporter under the control of the Rosa26 (R26R) promoter (Figure 4A). Animals initially received a 3-week pulse, followed by a long 14-week chase (Figure 4B). Any cells that expressed TERT during the doxycycline pulse, as well as their progeny after the chase period, would therefore express luciferase at the conclusion of the study, which was visualized via small animal imaging with an IVIS-CT. Luciferase signal was localized to brain regions including the olfactory bulb and olfactory epithelium (Figures 4C and 4D), as well as the lateral ventricles (LVs) (Figure 4E). Given the longer duration of this study, these signals may represent a combination of TERT cell proliferation, migration, and differentiation, but also confirmed the presence and contribution of TERT-expressing cells to basal brain cell turnover. Also of note, the IVIS-CT images are very sensitive to luminescent signal and the brightness may over-represent the number of cells expressing the tag.

**Figure 4:**
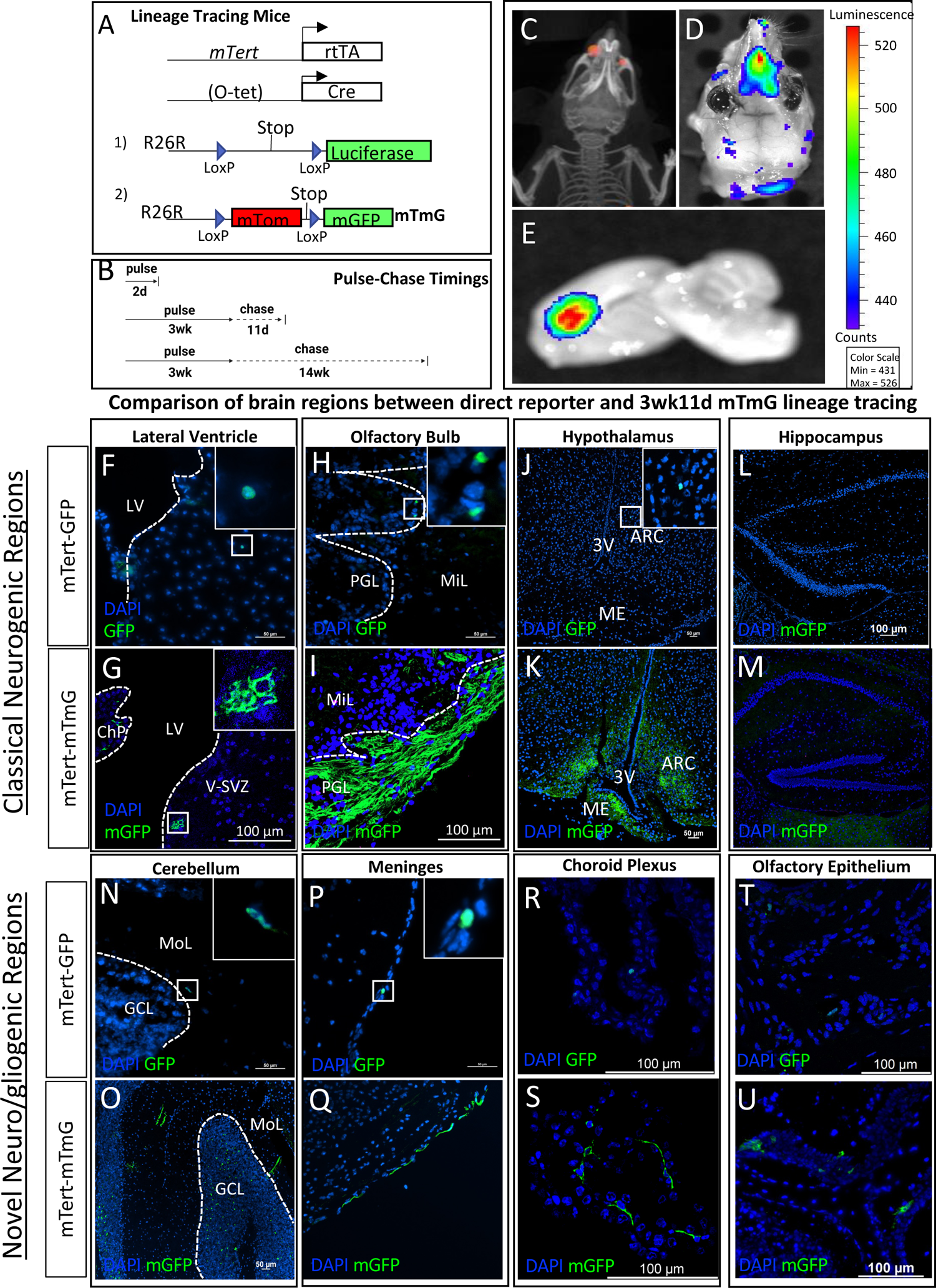
Lineage tracing revealed that mTert+ cells gave rise to a heterogeneous population of cells in multiple plastic regions of the brain. **(A)** Depiction of mTert-rtTA::oTet-Cre transgenic mouse models. 1) mTert-rtTA::oTet-Cre::LSL-Luciferase, 2) mTert-rtTA::oTet-Cre::R26R(mTmG). **(B)** Illustration of major pulse-chase timings utilized across both lineage-tracing mouse lines. **(C-E)** IVIS-CT of mTert-rtTA::TetO-Cre::Rosa-Luciferase lineage tracing mouse after a 3-week pulse, 14-week chase showed luciferase expression through the mouse head (C), the dorsal view through the skull (D), and the sagittal view of the brain (E) (*N* = 2 mice). The luminescence scale is shown to the right. **(F-M)** Representative images of TERT+ cells from immunofluorescent analysis across classical neuro/gliogenic brain regions in direct reporter mice versus mTert-mTmG mice after a 3-week, 11-day pulse-chase. Images taken of mTert-GFP brains in the LV (F), OB (H), hypothalamus (J), and hippocampus (L) (*N* = 2-11 of 11 mice showed this signal [*N* = 2-5 of 5 males, 0-6 of 6 females], *n* = 4-20 sections per brain analyzed). Representative images of adult mTmG lineage tracing mice after a 3-week, 11-day chase in the LV (G), OB (I), hypothalamus (K), and hippocampus (M); *N* = 6-10 of 10 mice showed this signal; *N =* 3-4 of 4 males, 3-6 of 6 females, *n* = 1-23 sections per brain analyzed). **(N-U)** Immunofluorescent analysis across novel neuro/gliogenic regions in mTert-GFP (N,P,R,T; *N* = 2-6 of 6-10 mice showed this signal [*N*=1-3 of 3-5 males, 1-3 of 3-5 females], *n* = 8-12 sections per brain analyzed) and mTmG mice after a 3-week pulse, 11-day chase (O,Q,S,U; *N* = 8-11 of 11 mice showed this signal [*N* = 4 of 4 males, 4-7 of 7 females], *n* = 1-19 sections per brain analyzed). Scale bars are 100µm. Insets show indicated area at 3x digital zoom. LV: Lateral Ventricle, PGL: periglomerular layer, MiL: mitral layer, 3V: third ventricle.

### GFP-marked TERT progeny cells expand in number after lineage tracing, including in classical neuro- and gliogenic niches of the adult mouse brain

For the remainder of our work, we used the lineage tracing mice with a fluorescent tag, the mTert-rtTA::oTet-Cre::R26R-mTmG (*mTert*-mTmG) mice.^4^ In these animals, recombination after doxycycline administration resulted in loss of membrane tomato expression and activation of membrane (m)GFP expression through the doxycycline-inducible Tet-On system (Figure 4A). In these mice, the fluorophore is not only membranous, unlike the direct reporter GFP, but it is an enhanced GFP signal and provides a brighter marker for TERT traced cells. We utilized various pulse-chase paradigms to better understand large-scale changes occurring throughout major TERT+ brain regions identified previously in the direct reporter GFP mice and to capture cell proliferation, migration and differentiation: 1) a short 2-day pulse, 0-day chase and/or 5-day pulse, 0-day chase (to closely mimic the direct reporters), 2) a 3-week pulse, 11-day chase, which we used for most analyses, and 3) a long 3-week pulse, 14-week chase similar to the IVIS data (Figure 4B). The 2-day pulse, 0-day chase was the shortest possible time required for mGFP induction and was used to compare to direct reporter mice ^33^. The 3-week pulse, 11-day chase was chosen because adult-born neurons are created in approximately 2-4 weeks in the SGZ, V-SVZ, and hypothalamus.^34–38^

Lineage tracing over the 3-week pulse, 11-day chase period revealed expansion of *mTert*-driven mGFP signal throughout various brain regions when compared to *mTert-*GFP direct reporter mice (Figures 4F-U). Immunostaining of *mTert*-GFP brains revealed TERT+ cells in proximity to the walls of the LV (Figure 4F). The expansion of this signal in the LV after a 3-week pulse, 11-day chase in *mTert-*mTmG animals appears to be minimal (Figure 4G). It is possible that TERT+ cells in the V-SVZ do not often give rise to V-SVZ or RMS cells under basal conditions, but instead respond to injury, metabolic, or other plasticity promoting cues, or that TERT+ qASCs are not a significant contributor to turnover of this niche.

In the OB of *mTert-*GFP mice, TERT+ cells were identified in the periglomerular layer (PGL; Figure 4H), mitral layer (MiL; Figure S4A), and granule cell layer (GCL; Figure S4A). Surprisingly high mGFP+ cell numbers were observed in the PGL and the olfactory nerve layer (ONL) of the OB after a 3-week, 11-day pulse-chase, indicating proliferation and differentiation activity of the qASCs in this niche (Figure 4I). Traditional V-SVZ neurogenesis leads to the formation of adult-born neurons via migration of TACs and neuroblasts through the RMS into the GCL of the OB, not the PGL and ONL. Progenitors and immature neurons that migrate to the OB via the RMS will integrate into the GCL more often than the PGL and ONL.^39^ In accordance with the low number of Tert-GFP+ and lineage traced mGFP+ cells in the V-SVZ (Fig. 4F-G), we also observed low numbers of mGFP+ cells within the GCL. These mGFP+ cells did not show a neuronal phenotype, indicating a possibly gliogenic pathway within or progenitor migration to the OB (Figure S4B). Although lineage tracing revealed mGFP+ cells within the RMS (Figure S4C), we never observed lineage traced cells with a neuronal morphology within the GCL that is typical of traditional V-SVZ-derived neurogenesis (Figure S4B). Instead, mGFP+ cells were found at low numbers in the mitral layer of the OB with similar morphology to the mGFP+ cells in the GCL (Figure S4D). It is therefore likely that adult neurogenesis from TERT+ stem cells in the olfactory epithelium (OE), as observed by luciferase signal after lineage tracing (Figures 4C and 4D), fueled the expansion of mGFP expression in the PGL and ONL.^39^

In the hypothalamus, adult neurogenesis regulates energy balance and metabolism via the development of new adult-born proopiomelanocortin (POMC)+ and neuropeptide Y (NPY)+ neurons, as well as hypothalamic glial cell differentiation.^35^ Metabolic interventions such as high fat diet can increase neurogenesis in the median eminence (ME) of female mice, while decreasing neurogenesis in the arcuate nucleus (ARC) of both male and female mice.^40^ Additionally, treatment with ciliary neurotrophic factor (CNTF), which causes weight loss in obese rodents and humans, increases cell proliferation in feeding centers of the murine hypothalamus ^41^. Weight loss in mice is reversed via coadministration of the mitotic inhibitor cytosine-beta-d-arabinofuranoside (Ara-C), indicating that hypothalamic adult neurogenesis is crucial for whole body energy balance.^35, 40^ Our immunofluorescent analysis revealed that the hypothalamus contained TERT+ cells in *mTert*-GFP direct reporter animals (Figure 4J), including cells nearby to the tanycyte region of the third ventricle (3V; Figure S4E). Tanycytes are radial glial-like cells that line the ventricles and are implicated in glucose-sensing and stem cell-like behavior.^42^ In the hypothalamus we also identified TERT+ cells in the ARC (Figure 4J). Compared to direct reporter mice, more GFP+ cells in the ARC and ME of the hypothalamus were present after lineage-tracing, suggesting TERT+ qASC contribution to tissue turnover in this niche (Figure 4K). Interestingly, in the hippocampal DG, no TERT+ cells were ever identified (Figure 4L), even after a 3-week pulse,11-day chase lineage tracing period (Figure 4M).

### TERT+ stem cells expand in number following lineage tracing

TERT+ cells were also identified in brain regions where adult neuroplastic potential has been observed but remains poorly understood. These regions, which included the cerebellum, meninges, and ChP, have been reported to contain proliferative stem cells with the ability to differentiate.^25, 43–45^ While TERT+ cells were sparse within the cerebellum in direct reporter mice (Figure 4N), lineage tracing resulted in a large-scale expansion of mGFP signal throughout various cell types and regions of the cerebellum that had not been observed in the *mTert-*GFP direct reporter animals (Figures 4N and 4O). We confirmed traced cell types in the cerebellum that had the morphological identity of Bergmann glia (Figure S4F), granule cells (Figure S4G), basket cells (Figure S4H), and Purkinje neurons (Figure S4I). TERT-traced cells in the cerebellum possessed clear morphological identity of mature cell types indicating that TERT+ cells give rise to mature adult-born cell types in the basal state of the adult mouse brain.

The meninges also harbored TERT+ cells in *mTert*-GFP mice (Figure 4P). Lineage tracing subsequently revealed mGFP+ expression increased compared to direct reporter expression in the meninges (Figure 4Q). These patterns were observed within the ChP as well (Figures 4R and 4S). The meninges and ChP brain regions sit at the blood-brain barrier and blood-cerebrospinal fluid (CSF) barrier interfaces and play a major role in immune cell trafficking^46^. The ChP produces CSF, which in turn coordinates neurogenesis via the circulation of growth factors throughout the ventricular regions of the brain, including the lateral and third ventricles, and hypothalamus.^47, 48^ These data indicate that TERT+ cells have the potential to give rise to mature adult-born cell types throughout non-neuronal and non-glial niches of the adult mouse brain. *mTert*-GFP animals had low numbers of TERT+ cells in the OE (Figure 4T), and after a 3-week, 11-day lineage tracing experiment, mGFP+ cells were found at higher numbers throughout the OE compared to the direct reporter animals (Figure 4U), supporting the idea that newborn olfactory sensory neurons (OSNs) in this niche may be observed within the olfactory NL and PGL. Taken together, TERT+ cells give rise to adult-born cells in most, but not all, classical and newly discovered neurogenic brain regions.

### Longer lineage tracing times increased expansion of TERT+ progeny

Expansion of the mGFP+ cell labeling over increasing pulse-chase times was observed in the OB, from the 2-day pulse, 0-day chase to the 3-week pulse, 14-week chase. It is important to note that these changes occurred primarily in the ONL and PGL (Figures 5A-C). The fact that TERT+ cells exist within the OB in direct reporter animals indicates the possibility of TERT+ cell proliferation within the OB instead of primarily via migration from the SVZ via the RMS, however, the extent of GFP expression after a pulse-chase in the PGL and NL implies that this occurs in combination with direct neurogenesis in the OE. mGFP+ cell expansion was also identified in the cerebellum between the 2-day pulse, 0-day chase and 3-week pulse, 14-week chase (Figures 5D-E), but fluorescent intensity was reduced in the 3-week, 14-week pulse-chased cerebellum (Figure 5E), indicating that TERT-traced cell expansion within this niche may be a transient, short-term phenomenon. Within the hypothalamic ARC a trend was observed towards increased fluorescent intensity between 2-day pulse, 0-day chase and the other pulse chase times (Figures 5F-G). The thalamus also showed a trend towards increased mGFP signal intensity between animals treated with a 2-day pulse, 0-day chase and 3-week pulse, 11-day chase (Figures 5H-I). Overall, across the three pulse-chase paradigms we rarely observed mGFP traced cells in the V-SVZ and RMS (Figures S5A-5B) despite these being classical adult stem cell niches in the mouse brain.

**Figure 5:**
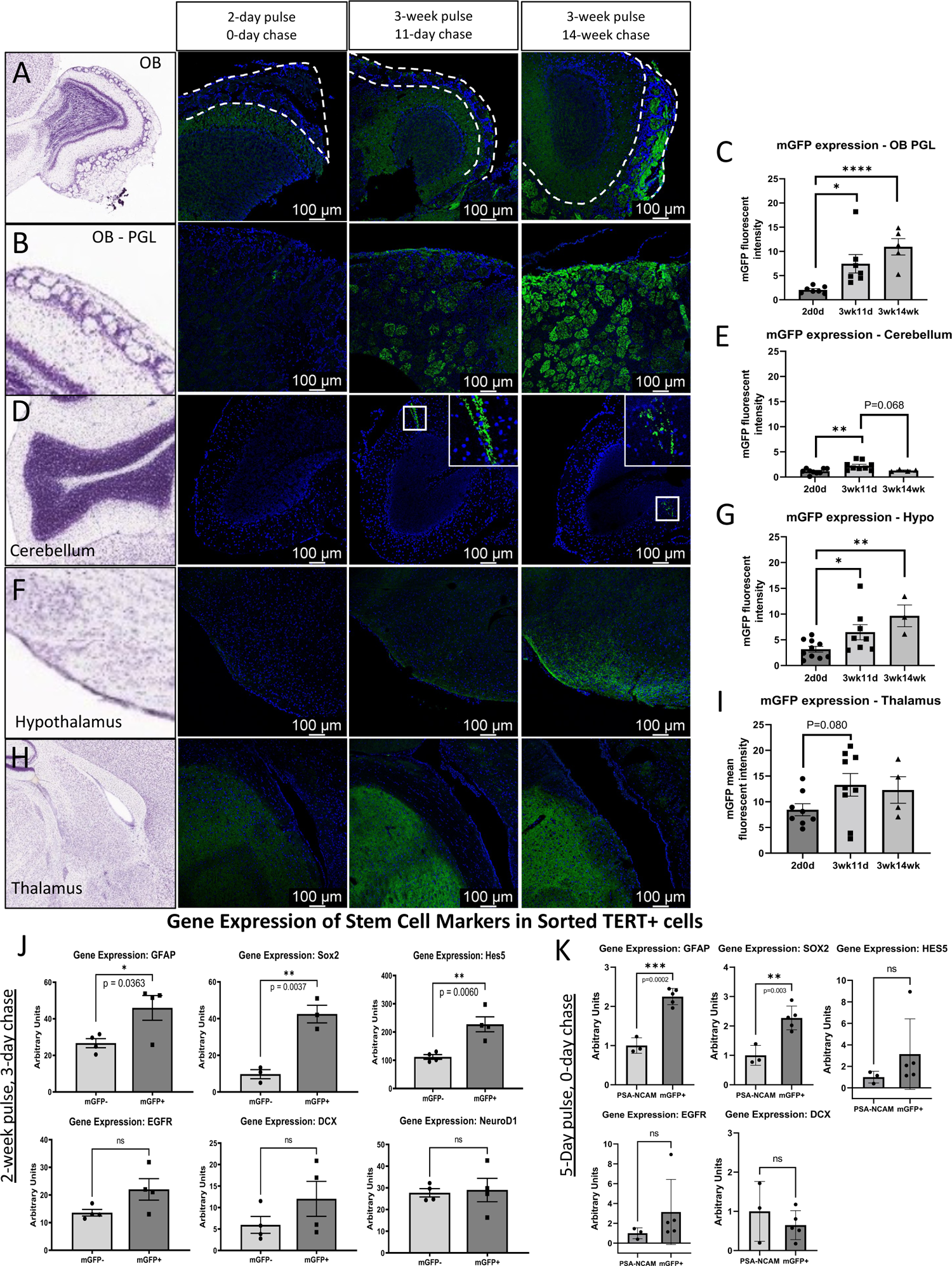
Lineage tracing revealed numerous brain regions with high basal plasticity and differentiation of TERT+ precursors that increased with pulse-chase time. **(A-I)** Representative images of each of the OB (A), OB PGL (B), cerebellum (D), hypothalamus (F), and thalamus (H). Representative images of sagittal-view brain regions of interest were obtained from the Allen Brain Atlas. Quantification of mean fluorescent intensity of mGFP expression (± SEM) per mouse across lineage tracing in the OB PGL (C), cerebellum (E), hypothalamus (G), and thalamus (I). mTert-mTmG mouse brains were immunostained following a: 2-day pulse, 0-day chase (*N* = 5 males, 5 females); 3-week pulse, 11-day chase (*N* = 4 males, 8 females); or 3-week pulse, 14-week chase (*N* = 6 females). For each brain region imaged, *N* = 4-10 mice, *n* = 1 section per brain. Dots represent mean fluorescent intensity of individual animals (*N* = 4-10; \**P* < 0.05, \*\**P* < 0.01, \*\*\*\**P* < 0.001). All scale bars are 100µm. All images taken at the same sagittal depth. Dotted lines indicate the PGL identified in the OB. **(J)** qRT-PCR analysis of mGFP+ traced cells sorted via MACS and PSA-NCAM+ cells sorted via FACS. Expression of SOX2, GFAP, EGFR, DCX and HES5 was normalized by Cyclophilin expression of the PSA-NCAM+ cells (N =5; 2 males and 03 females for mGFP, N=3; 2 males and 1 female for PSA-NCAM sorted cells, \*\**P* < 0.01, \*\*\*\**P* < 0.001). **(K)** qRT-PCR analysis of transcripts from mGFP+ traced cells sorted via FACS after a 2-week pulse, 3-day chase for markers of stem cells and activation and neuronal differentiation. *N* = 4 males pooled to 2 samples, *N* = 2 females.

Intracerebroventricular (*i.c.v.*) treatment with the mitotic inhibitor Ara-C resulted in a reduction of DCX+ cells along the lateral ventricle wall/V-SVZ (Figures S5C-E), as expected, but did not result in changes to the fluorescent intensity of lineage-traced mGFP expression in cells in the LV ChP (Figure S5F). This makes sense since we observed TERT+ cells as mainly quiescent, and AraC targets only mitotic and dividing cells, thus sparing the TERT+ quiescent cells. However, Ara-C treatment did not significantly decrease mGFP expression in the OB, indicating that the mGFP signal within the OB likely does not originate from the V-SVZ (Figure S5G-I).

FACS analyses after a 2-week, 0-day pulse chase (thus, matching the direct reporter brains) revealed that mGFP cells accounted for 0.2% of the full adult mouse brain and these isolated cells formed neurospheres in culture, similar to what was observed in the *mTERT*-GFP direct reporter mice (Figure S5J). The percentage of mGFP cells in the adult brain increased to 3.8% after a 3-week, 11-day pulse-chase (Figure S5K), consistent with our immunostaining observations. These lineage-traced cells from a longer pulse-chase exhibited neurosphere formation as well (Figure S5J-K), indicating they contained progenitor cells and not just mature cell types.

After a relatively short 2-week pulse, 3-day chase, performed so that intermediate progenitor cell types resulting from TERT+ cells could be identified, GFP+ cells isolated via FACS from mTert-mTmG brains expressed significantly higher levels of mRNA transcripts for quiescent and activated stem cell markers (*GFAP, Sox2,* and *Hes5)* compared to all mGFP-negative cells from the same brains (Figure 5J). However, transcripts for markers of activated ASCs, immature neurons and mature neurons (*EGFR*, *DCX*, and *NeuroD1)* were not significantly different between mGFP+ and mGFP-cells at this pulse-chase timepoint. Additionally, we isolated mTERT+ cells after a 5-day chase, 0-day pulse through MACS sorting, a more gentle separation approach, to investigate the gene expression of several quiescent and activated ASC markers: SOX2, GFAP, HES5, DCX and EGFR (Figure 5K). This time we utilized PSA-NCAM+ cells as a negative control, to represent a committed and/or differentiated neuronal population that does not contain stem cells, instead of using every GFP-negative cell in the brain which likely included dozens of cell types. These results indicated increased mRNA levels for GFAP and SOX2 in the mGFP+ cells when compared to the PSA-NCAM+ cells (p=0.0002 and p=0.003, respectively).

To further characterize the quiescent vs activated state of TERT+ stem cells, we compared the metabolic activity of sorted cells in culture through a Seahorse respirometry assay that quantified oxygen consumption rate (OCR) and extracellular acidification rate (ECAR). After 5 days in culture the TERT+ cells displayed very low levels of OCR and ECAR (ranging from 0.5-1.5 pmol/min and 1-4 mpH/min, respectively). By contrast, after formation of neurospheres stimulated by the presence of EGFR and FGF in the media for 15 days, these cells were more metabolically active exhibiting 10-40 pmol/min OCR and 2-10 mpH/min ECAR levels (Figure S6A-D).

### TERT+ cells gave rise to both immature and mature neuronal cell types following lineage tracing

In order to identify the cell types arising from lineage-traced TERT+ populations across the adult mouse brain, we stained mGFP+ cells from *mTert-*mTmG lineage traced mice with markers of activated ASCs, TACS, neuroblasts, and neurons. These data are represented in Figure 6A. The neuronal differentiation pathway is again shown in Figure 6B for reference. In the V-SVZ, the activation marker EGFR never co-localized to lineage traced mGFP+ cells after a 2-day, 0-day pulse-chase, nor the 3-week, 11-day pulse-chase, and only co-localized rarely with mGFP+ cells from 3-week, 14-week lineage-traced *mTert*-mTmG animals (Figure 6C). While these data indicate that TERT+ cells often do not become activated, there is likely a transient activation period which is technically challenging to capture in these types of lineage tracing experiments.

**Figure 6.**
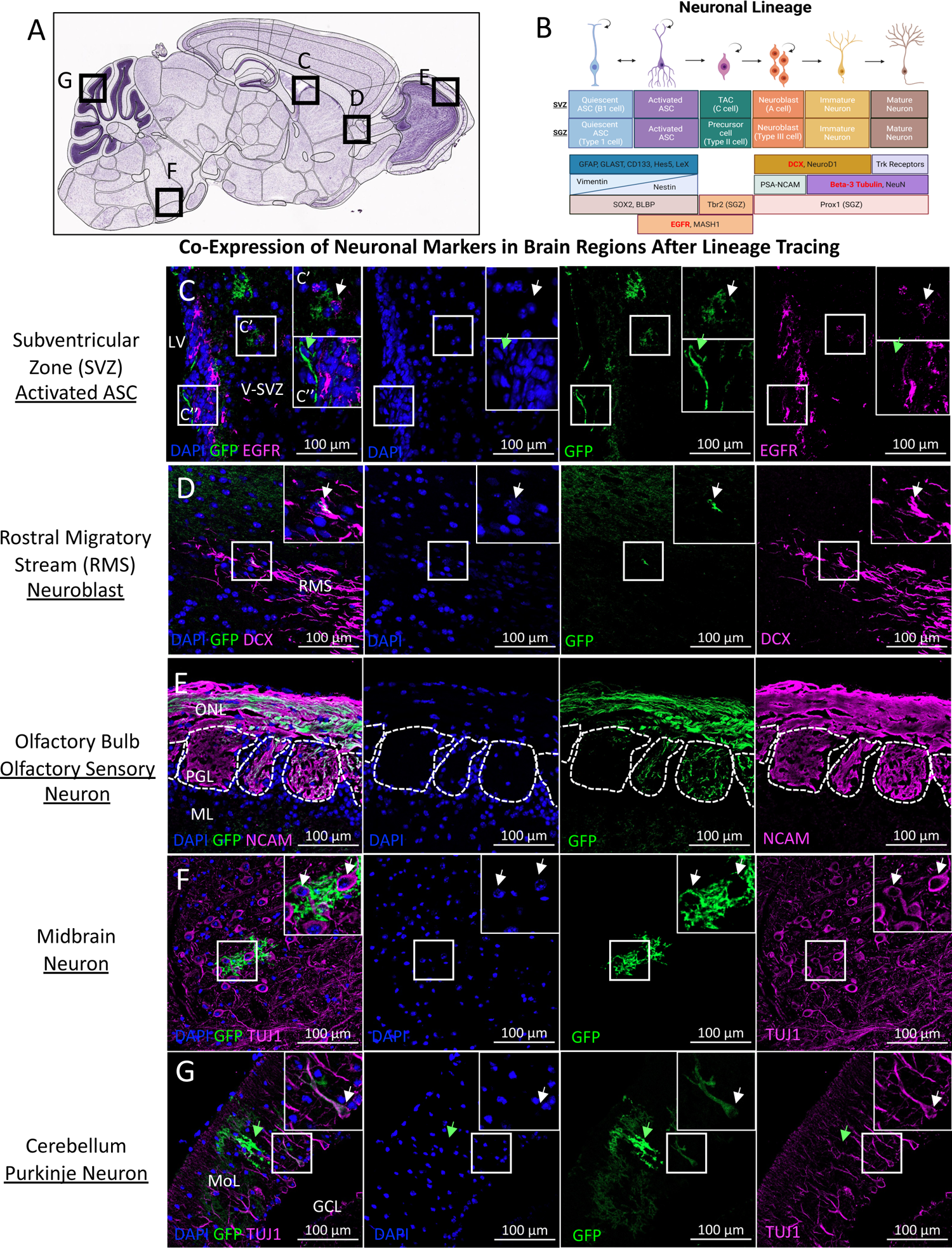
Lineage tracing of TERT+ cells revealed immature and mature neuronal cell types throughout the adult mouse brain, confirming the multipotency of TERT+ ASCs. (A) Sagittal mouse brain indicating the region each image was captured C-G (Allen Brain Atlas). (B) Schematic depicting neuronal lineage markers; quiescent stem cell to mature neuron. Markers with red text were stained for in this image. (C) Co-staining of lineage tracing mice after a 3-week, 14-week pulse-chase with the activated marker (EGFR). Both mGFP+EGFR+ (C’, white arrow) and mGFP+EGFR-(C’’, green arrow) cells were identified. No other mGFP+ cells expressed EGFR in the V-SVZ (*N* = 5 males, 5 females with 2-day, 0-day pulse-chase, *n* = 2 sections per brain; *N* = 4 males, 4 females with 3-week, 11-day pulse-chase, *n* = 2 sections per brain; *N* = 6 females with 3-week, 14-week pulse-chase, *n* = 2 sections per brain). **(D)** Immunostaining of mTert-mTmG brains after a 3-week, 11-day pulse-chase stained with the neuroblast marker DCX in the RMS (seen in *N* = 1 of 4 mice [*N* = 1 of 2 males, 0 of 2 females], *n* = 2 sections per brain analyzed) **(E)** Representative image of co-immunostaining of OB stained with the olfactory sensory neuron marker NCAM after 3-week, 11-day pulse-chase (seen in *N* = 4 of 5 mice [*N* = 2 of 2 males, 2 of 3 females], *n* = 2 sections per brain). **(F-G)** Representative images of co-immunostaining with the mature neuron marker TUJ1 in the brainstem (F; *N* = 2 of 4 mice [*N* = 1 of 2 males, 1 of 2 females]) and cerebellum (G; *N* = 3 of 4 mice [*N* = 2 of 2 males, 1 of 2 females]) after a 3-week, 11-day pulse chase. Scale bars are 100µm. Co-stained cells are indicated with white arrows. Cells that express mGFP only are indicated with green arrows. Insets show 4x digital magnification of indicated area. RMS: rostral migratory stream, MoL: molecular layer, GCL: granule cell layer, V-SVZ: ventricular-subventricular zone.

After a 3-week, 11-day lineage trace, mGFP+ cells in the ventricular lining and V-SVZ were negative for Sox2 (Figure S7A). Interestingly, mGFP+Sox2+ cells were found in the midbrain (Figure S7B) and in the RMS (Figures 6D and S4C). A subpopulation of mGFP+ traced cells in the RMS were DCX+ neuroblasts (Figure 6D). mGFP+ cells found in proximity to the RMS were DCX- and lacked a the neuroblast phenotype (Figure S7C), indicating they may not be migrating neuronal precursors. The activation marker EGFR was also co-expressed by lineage-traced mGFP+ cells in the brainstem (Figure S7D). Although no mGFP+ cells were found in the SGZ of the hippocampus after a 2-day, 0-day pulse-chase, nor following the 3-week, 11-day pulse-chase, we did observe a single mGFP+ cell cluster in the DG of the hippocampus after the very long 3-week, 14-week pulse-chase (Figure S7E). This cluster of cells in the DG was EGFR+ (Figure S7E), and these cells may have migrated here in a rare event, given the lack of GFP expression in the hippocampus in any other mice across 20+ cohorts of this project thus far. Taken together, mGFP+ cells in the V-SVZ, RMS, and DG have the capacity to express markers of ASCs or progenitor cell types, indicating a cell type that has the potential to become activated and differentiate in neurogenic brain regions, but TERT+ expression in these classical niches is far lower than in non-classical niches of the brain.

Lineage-traced cells co-expressed the mature neuron axonal marker NCAM in the olfactory bulb (Figure 6E). NCAM is expressed only in the ONL and PGL and marks all PGL compartments due to expression of NCAM in olfactory sensory neurons.^49^ mGFP signal was highly expressed in the ONL but did not fill all PGL compartments after either a 3-week, 11-day, nor after the longest 3-week, 14-week pulse-chase (Figure 6E). These mGFP+ PGL axons also expressed vimentin, which is known to be expressed by OSNs within the PGL and NL (Figure S7G).^50^ Unexpectedly, about half of female animals showed little to no mGFP signal in the ONL and PGL after a 3-week, 11-day pulse-chase, whereas all of the males had bright signal (Figure S7F). This was a rare display of sexual dimorphism in these mouse models, whereas in other assessments we could not distinguish any sex difference in TERT cell numbers, anatomical localization, or lineage tracing cell fates. The mature neuron marker TUJ1 was found expressed by mGFP+ cells in the midbrain, identifying these cells as mature neurons (Figures 6F). mGFP+ cells in the cerebellum also expressed TUJ1 and showed morphology indicative of mature neurons (Figure 6G). Taken together, it is clear that TERT+ cells can give rise to a variety of neuronal cell types throughout the brain, which we confirmed by co-staining with mature cell markers, in both classic and non-classic neurogenic regions. However, consistent with other reports in the literature, we also find that the majority of adult brain basal tissue regeneration is due to non-neuronal cell types.

### TERT+ cells gave rise to numerous non-neuronal cell types, including glia, following lineage tracing

After a 3-week, 11-day pulse-chase, mGFP+NG2+ cells were identified in the cerebellum (Figure 7C). These cells were therefore likely either GPCs or oligodendrocyte-restricted progenitors, whereas NG2+ cells in the meninges were not mGFP+ (Figure S8A). Lineage-tracing also revealed mGFP+OLIG2+ oligodendrocyte-lineage cells in a variety of brain regions, including the cerebellum, RMS, midbrain, and islands of Calleja (IoC; Figures 7D-7E and S8B-C). mGFP+GFAP+ cells in the astrocyte-lineage were observed directly adjacent to and within the ventricular lining in the fourth ventricle (4V; Figure S8D), and in dense regions of the ventral hypothalamus (Figure 7F). Importantly, lineage-traced mGFP+ cells were identified lining all the ventricles of the adult mouse brain (4^th^ ventricle shown in Figure 7G), and may be positioned near the CSF barrier in order to gain access to circulating growth factors – a common feature of stem cells in various tissues which may form niches near the blood vasculature, for example.

**Figure 7.**
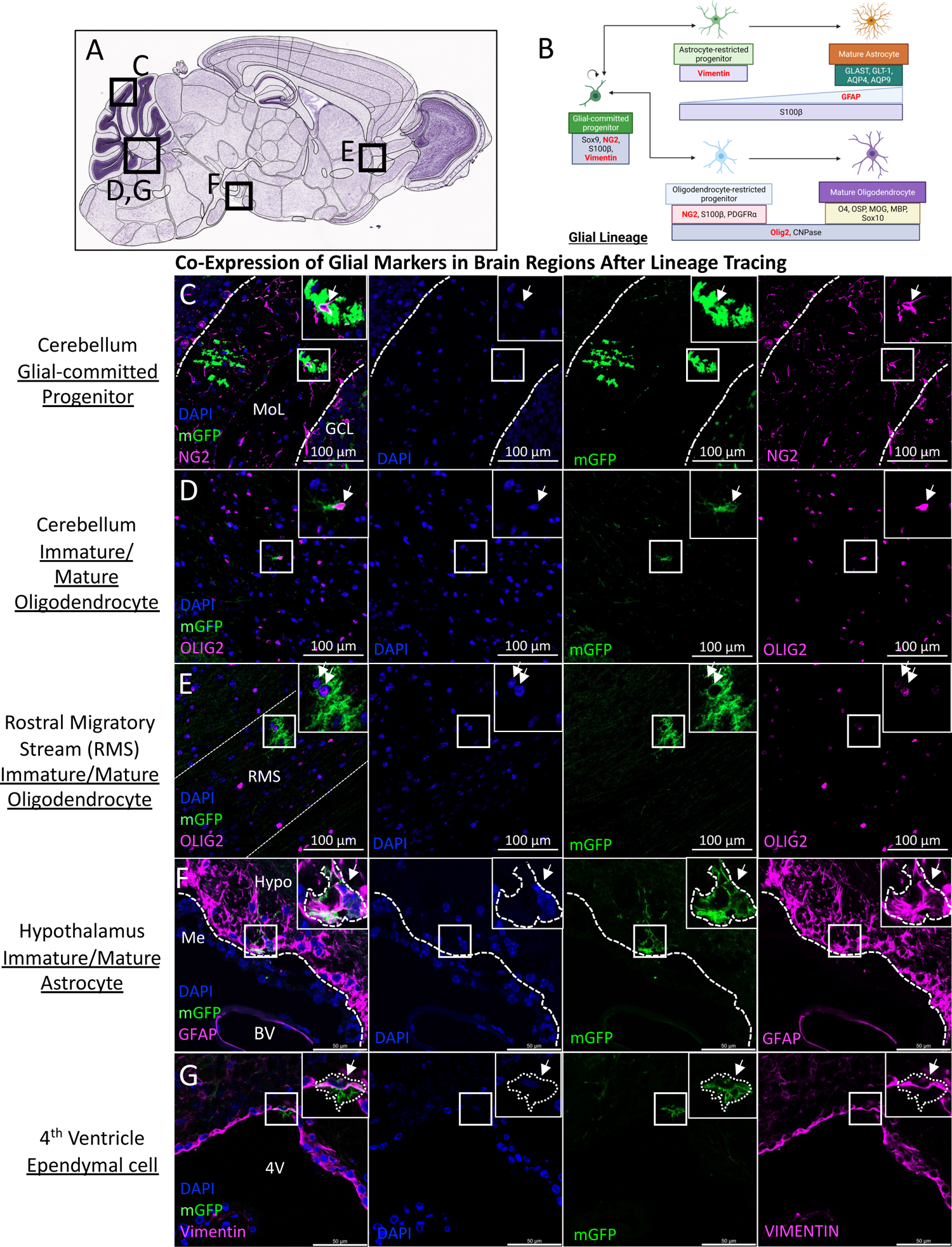
Lineage tracing of TERT+ cells revealed immature and mature glial cell types throughout the adult mouse brain, confirming the multipotency of TERT+ ASCs. **(A)** Sagittal mouse brain indicating areas of images C-G (Allen Brain Atlas). **(B)** Schematic depicting neuronal lineage markers of glial-committed progenitors through mature astrocytes and oligodendrocytes. Red text indicates markers stained in this figure. **(C-G)** Co-staining of 3-week 11-day pulse-chased lineage tracing mice with the glial progenitor marker NG2 (C), the oligodendrocyte marker Olig2 (D-E), the hypothalamus (F), and the 4th ventricle (G; *N* = 4-5 of 5 males, 5-6 of 6 females showed this signal, *n* = 2 sections per brain analyzed). Scale bars are 100µm. White arrows indicate cells that co-express markers of interest. Insets show 2x digital zoom of indicated area. RMS: rostral migratory stream, MoL: molecular layer, GCL: granule cell layer, BV: blood vessel. 4V: 4th Ventricle

### Brain clearing identified lineage-traced TERT+ cells and their progeny throughout the adult mouse brain

To gain a whole-organ perspective of TERT+ lineage traced cells throughout the brain, we performed brain clearing on mTert-mTmG brains after a 3-week, 11-day pulse-chase, followed by immunostaining to boost the GFP signal (which can often be lost due to clearing-induced quenching of endogenous fluorophores). Brain clearing via iDISCO allowed for thicker segments of the adult mouse brain to be stained and imaged via confocal microscopy in the sagittal plane.^23^ We observed GFP+ signal in the OB-PGL (Figure 8A), cortex (Figure 8B), cerebellum (Figure 8C), brainstem (Figure 8D), ChP (Figure 8E), and meninges (Figure 8F). These patterns are similar to what we observed in thin-section immunostaining from Figures 4-7. Interestingly, high cortex signal was observed in lineage traced animals (Figure 8B) when viewed in this manner, which allows a maximum projection of cells expressing GFP across the depth of the tissue (and thus reveals cells that may appear insignificant in thin sections).

**Figure 8.**
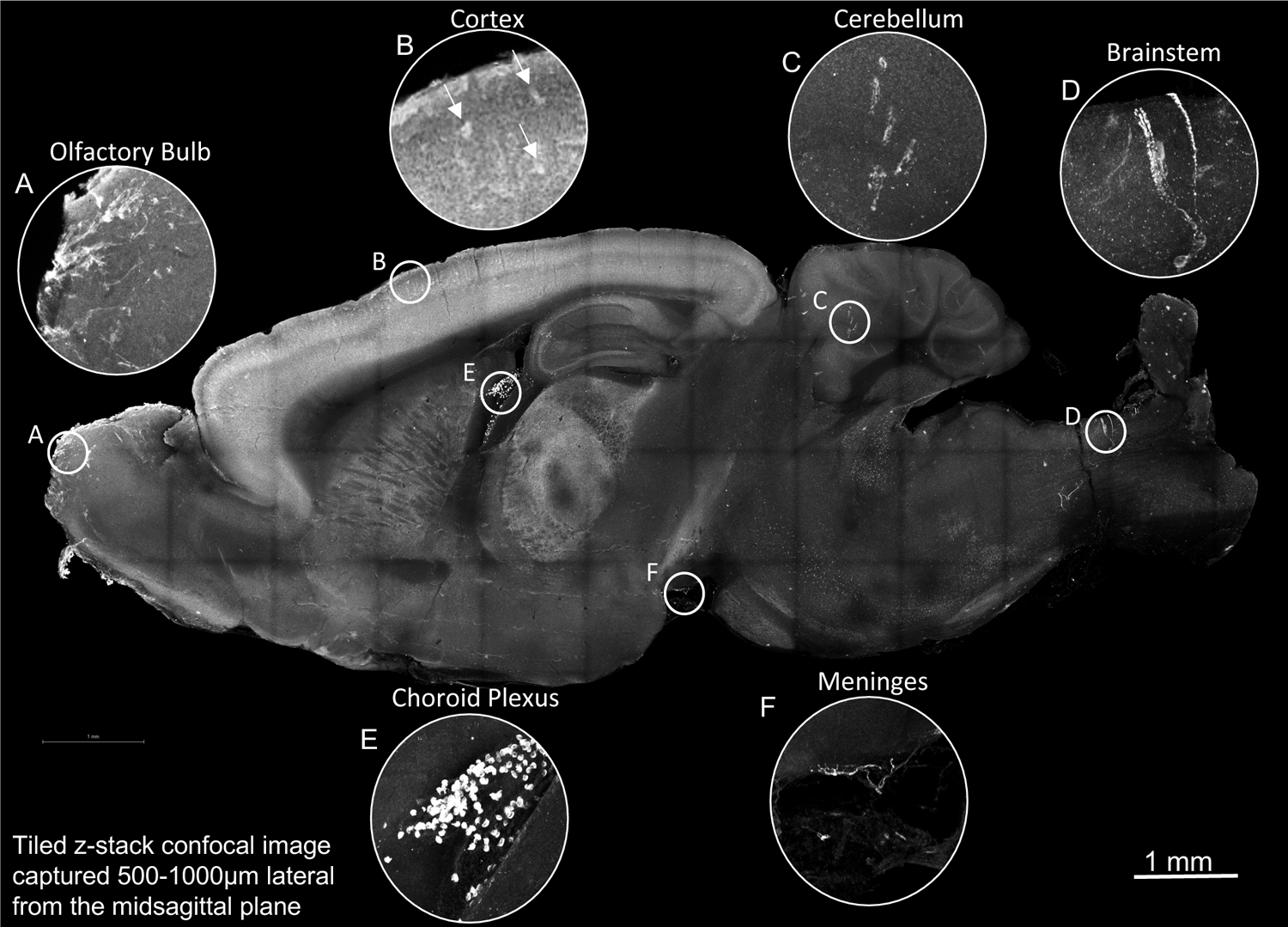
Lineage tracing identified TERT+ cells and their progeny throughout the adult mouse brain. **(A-F)** Representative immunostaining for GFP within a 500µm cleared sagittal brain section of an adult mTmG mouse brain after a 3-week, 11-day pulse-chase showed GFP+ signal (grey) in the A) OB PGL, B) LV ChP, C) cerebellum, D) brainstem, E) meninges, and F) cortex (*N* = 3 mice, *n* = 2 sections per brain). White arrows indicate GFP+ cells in 8F. Insets show 5x digital magnification of indicated areas. Images acquired on a Leica Stellaris confocal microscope. Images were captured at 10x, tiled and Z-maximum intensity projected to create an intact rendering of 500-1000µm from the midsagittal plane.

## Discussion

Although adult neurogenesis was first discovered in avian brains in 1984 ^51^ and murine brains in 1992,^52, 53^ the first evidence of human adult neurogenesis was as recent as 1998.^54^ The current understanding is that stem cells capable of developing into new mature neurons in the adult mammalian brain are restricted to certain neurogenic niches, and likely lose this neurogenic capacity as we age. Recently, the view that adult neurogenesis only occurs in the V-SVZ and SGZ has been challenged by the identification of adult stem cells that can give rise to neurons and glia in numerous other brain regions in rodents, such as the ChP ^25^, meninges ^45^, cerebellum ^43^, and more.^2^ Here, we report the identification of a novel, rare, largely quiescent, and multipotent population of TERT+ ASCs in numerous anatomical niches of the adult murine brain. These TERT+ ASCs are somewhat distinct from GFAP+/Nestin+/Sox2+ cells, often termed quiescent adult neural stem cells (qANSCs), which are found within the SGZ or V-SVZ. While GFAP+/Sox2+/Nestin+ cells in the adult mouse brain are restricted to differentiation to neurons, astrocytes, and oligodendrocytes, we have demonstrated that TERT+ ASCs are truly multipotent and can differentiate into various subtypes of neurons, astrocytes, oligodendrocytes, ependymal cells, and ChP epithelial cells. TERT+ ASCs are also less restricted anatomically than classical qANSCs, and we observe them in the basal state in numerous plastic brain regions including the V-SVZ, OB, and hypothalamus, but strikingly never in the hippocampus. These differences indicate the possibility that previously described ANSCs in the V-SVZ and SGZ are either a separate stem cell population than TERT+ ASCs, or a subset.

In the adult mammalian brain, the V-SVZ contains qANSCs known as Type B1 cells.^55^ These cells can become activated and asymmetrically divide into self-renewing B1 cells and transit amplifying cells (TACS), also known as Type C cells. TACS then further divide to produce neuroblasts (Type A cells), which migrate rostrally within the rostral migratory stream (RMS) to the olfactory bulb (OB), where they differentiate into mature neurons that integrate into the neural circuitry.^31^ In humans, V-SVZ-derived progenitors appear to migrate to the striatum instead of the OB.^56^ Within the SGZ, quiescent radial glial-like stem cells (Type 1 cells) produce intermediate progenitor cells (Type 2 cells) via symmetric or asymmetric cell division. These cells produce adult-born neuronal cells that express doublecortin (DCX). DCX+ cells then migrate into the granule cell layer where they differentiate into mature DG neurons.^57^ By contrast, in other reportedly neurogenic regions such as the hypothalamus and cortex, the cell types and mechanisms involved in adult neuro/gliogenesis are less well understood and potentially new stem cell types in the adult mouse brain have not yet been described.^2^

Despite the identification of molecular markers expressed by ANSCs in prior research, a specific and unique marker for multipotent ASCs across the adult mammalian brain has remained elusive^58^. Nestin, GFAP, GLAST, Sox2, and other markers have been proposed to mark slowly cycling multipotent stem cells, but each of these also marks multiple non-stem cell types throughout the brain (ex: Nestin marks endothelial and meningeal cell types, GFAP and GLAST mark mature glia, and Sox2 is widely expressed throughout the brain). TERT is an appealing candidate to mark qASCs, given the numerous prior studies validating TERT+ qASCs in multiple other tissues and organs.^3, 4, 6, 10, 59, 60^ We have now demonstrated that TERT represents a unique and specific marker of adult ASCs in the brain, as supported by the low frequency of TERT+ cells in direct reporter mice, the lack of proliferation of these cells in unstimulated conditions, and the ability of these cells to contribute to tissue turnover of mature numerous cell types as demonstrated in lineage tracing studies.

As presented, a large percentage of TERT+ cells (53%) in the adult *mTert*-GFP mouse brain express the immune cell marker CD45 (Figure S1). These are likely circulating lymphocytes since the majority of TERT+/CD45+ cells were identified in the ChP and meninges, brain regions with high immune cell infiltration. This is further supported by data demonstrating that human TERT (hTERT) is expressed by lymphocytes isolated from thymus, tonsil, and peripheral blood.^61^ hTERT expression is also found in microglia expressing CD68 in adult human brain, indicating that TERT expression occurs in activated microglia. ^62^ Therefore, TERT+CD45+ co-expressing cells may be infiltrating immune cells from the periphery via the bloodstream and glymphatic system, or represent activated resident microglia. These are therefore distinct from the TERT+ ASCs in the brain parenchyma as we describe. Previous studies by our colleagues, using the same TERT mouse models, have investigated the transdifferentiation ability of bone marrow derived TERT+/CD45+ cells and concluded that these cells do not transdifferentiate into stroma, epithelium, and endothelium in the endometrium.^63^ Thus, TERT+CD45+ cells likely represent a committed or mature cell type that retains TERT expression, instead of a plastic/stem/progenitor cell type capable of renewing adult tissues as we have investigated via lineage tracing. Additionally, since CD45+ lymphocytes are post-mitotic, they are unlikely to contribute in any meaningful way to the lineage tracing observations in this study. It remains a possibility that a subpopulation of TERT+ cells are CD45-low microglia, which could have mitotic properties.

TERT+ cells in the adult brain represent a potentially heterogeneous population due to their variable expression of Ki67, Nestin, and Sox2 as observed in TERT-GFP+ direct reporter cells. This may be because TERT+ cells are largely dormant, or quiescent, and can become activated in response to stimulation – thus shifting their gene expression profile. TERT was originally hypothesized to be expressed only by proliferative cells, since the telomerase enzyme (composed of TERT and a telomerase RNA component) prevents telomere shortening during chromosomal replication, thus allowing for continued cell division.^64^ However, in other adult murine tissues such as the intestine, where TERT+ stem cells are low in number and rarely express Ki67 (like our brain TERT+ cells), TERT clearly marks a population of adult tissue stem cells that are slowly cycling and quiescent. TERT may therefore carry out non-canonical functions beyond telomere extension in quiescent stem cells. Indeed, tight mitochondrial control of the balance between glycolysis and oxidative phosphorylation is required in stem cells, and TERT is understood to influence mitochondrial function distinct from its role in telomere extension.^65^ Additional non-canonical roles of TERT have been identified that may regulate the stem-ness of TERT+ cells, including regulating cell survival and differentiation capacity.^66, 67^ The discovery of a mostly non-proliferating pool of TERT+ stem cells capable of multipotent tissue regeneration in the intestine, bone marrow, liver, adipose, long bone, kidney, heart, and now brain, provides strong collective evidence for TERT as a marker of quiescent adult tissue stem cells.^3–6, 10–12, 59, 60^

The ability of TERT+ cells to form neurospheres *in vitro* further supports the identity of these cells as neural stem cells, as the formation of neurospheres is a characteristic response whereby stem cells can respond to cues such as the growth factors found in neurosphere culture media that activate the stem cells to proliferate and/or differentiate. As TERT+ cells appear to be quiescent *in vivo*, it is possible that the loss of GFP observed over time in our neurosphere cultures *in vitro* occurred due to activation by EGF and bFGF in the culture media, leading to a proliferative phenotype and loss of TERT expression. Interestingly, both TERT+/GFP+ sorted cells from *mTert*-GFP mice, and those that lost GFP expression in culture, incorporated EdU over 24 hours, indicating proliferation and cell division. While the expression of GFP would indicate that the neurospheres in culture retained TERT expression and quiescence, it remains possible that TERT expression was re-induced in cells, which had previously lost TERT through activation by the growth factors in the culture media. The re-expression of TERT in this case may have also been in response to telomere shortening via the process of cell division.

*mTert-*GFP direct reporter mice express GFP under the control of 4.4kb fragments of the mTert promoter region.^3^ These cells express high levels of Tert mRNA transcript, as well as variable levels of GFP.^3^ Analysis of TERT expression in this mouse model has been hindered by the lack of a validated TERT antibody. While the caveat remains that TERT-GFP+ cells could only be expressing the Tert mRNA and not translating the TERT protein, prior studies have confirmed that TERT-GFP+ cells exhibit telomerase activity, indicating the presence of functioning TERT protein within these cells.^3^ These TERT mouse models have also been utilized and validated in numerous tissue stem cell studies prior to our work in brain. 3-6,10,11,60

Although the V-SVZ and SGZ are classical neurogenic niches, there remains debate about how much of the observed neurogenic capacity in these rodent brain regions translates to the adult human brain, as there are substantial discrepancies reported between human and mouse adult neurogenesis in both the V-SVZ and SGZ.^68^ It is not yet fully understood where adult-born neurons from the human V-SVZ transit/migrate, as observations of the human RMS have never indicated migration of neuroblasts to the human OB. However, some research indicated that progeny of human V-SVZ ANSCs are destined for the striatum rather than the OB, which is much smaller and less utilized in humans than mice.^56^ In the human hippocampus, neurogenesis analyses have relied heavily on the presence of immature neurons (or committed neuroblasts), which are proliferative or label retaining cells.^54, 69–73^ Whether these immature neurons are derived from a population of uncommitted and multipotent stem cells within the niche, were created earlier in life to generate a pool of committed progenitors, or migrated to the hippocampus from other brain regions, is currently unresolved.

Additionally, depending on the method of analysis, adult human brains demonstrated rare or nonexistent hippocampal neurogenesis.^74–76^ For this reason, the presence of adult hippocampal neurogenesis in the DG after a few months into adult human life is still contested.^72, 76–81^ Our data support the hypothesis that the hippocampus houses committed progenitors, and not multipotent stem cells, since we never see TERT or its progeny in the hippocampus. Finally, it must be noted that while Tert mRNA expression and telomerase activity in the adult brain has been observed in mice ^82^, in humans, TERT *<u>expression</u>* has been observed in hippocampal neurons and activated microglia ^62, 83, 84^, but there has not yet been any evidence of human telomerase *<u>activity</u>* in the adult brain.^84, 85^ In fact, human telomerase activity is decreased in the brain to undetectable levels after the 16^th^ week of gestation.^85^ This may be due to limited tools to detect TERT protein in human brain samples, or the fact that adult human brains have TERT+ qASCs at very low frequency, as we see in mouse, making them difficult to detect.

TERT+ cells, and lineage-traced expansion of TERT+ cells and their progeny, were identified throughout the adult mouse brain, including outside of the classical neuro/gliogenic niches, but within niches with prior evidence of adult neuro/gliogenesis. Lineage tracing revealed an increased number of GFP+ cells (meaning they derived from TERT+ precursors) compared to TERT direct reporter animals, across brain regions that included the meninges, ChP, cerebellum, brainstem, and thalamus. The presence of a rare population of TERT+ ASCs in these regions that gave rise to large numbers of multiple cell types after a pulse-chase period clearly demonstrated that adult neuro/gliogenesis occurs in a far less restricted manner than previously believed, at least derived from TERT+ precursors. During development, neuroblasts migrate from the early postnatal V-SVZ to the medial prefrontal cortex, striatum, nucleus accumbens, and IoC. This process becomes restricted to only migration to the OB through the RMS during adulthood in rodents, but newborn neurons have been reported in the IoC in adult mice and rats, indicating the presence of alternative migratory pathways. While no TERT+ cells were found within the IoC in *mTert-*GFP animals, the presence of high mGFP signal in this region after lineage tracing in adult mice indicates migration of TERT+ progeny may also occur via the IoC, or other yet-to-be identified migratory pathways that may be predominantly used by TERT+ progeny.

In order for long-lived quiescent adult neural stem cells (qANSCs) or tissue adult stem cells (ASCs) to survive into adulthood and maintain a dormant reservoir of cells capable of tissue turnover and regeneration, their quiescence must be preserved over long periods. Quiescence is a state of reversible non-proliferation maintained by a transient gene expression program that promotes dormancy, such as reduced expression of cell cycle and DNA replication genes, as well as genes for mitochondrial function.^87^ On the other hand, mechanisms that prevent replicative senescence, or the irreversible cell cycle arrest and loss of proliferation that is characterized by impaired telomere activity^88^, include metabolic reliance on glycolysis rather than oxidative phosphorylation,^89^ expression of metabolic sensors such as forkhead box O (FOXO) to protect against reactive oxygen species,^87^ and maintenance of telomere length via activity of the enzyme telomerase.^90^

Based on our data, we hypothesize that maintenance of a TERT+ population of ASCs in the brain is necessary to retain neurogenic and gliogenic potential throughout adulthood. Dysfunction in the maintenance of this population may play a role in the cognitive deficits seen in aging and neurodegenerative diseases. For example, TERT KO animals show decreased V-SVZ neurogenesis and stem cell numbers in adults, implicating TERT in stem cell renewal and neurogenic niche function.^20^ Consistent with this hypothesis, analysis of telomerase activity in the V-SVZ after injury shows that TERT+ cells may be injury-responsive.^15^ In other adult tissues, TERT+ cells respond to injury by significantly increasing their proliferation and turnover. For this reason, it is possible that these cells may proliferate slowly under homeostatic conditions, even in brain regions where other highly proliferative cell types reside.^10, 59^ The fact that telomerase activity in the V-SVZ is induced after injury may explain why low numbers of TERT+ cells and lineage-traced cells were observed in the V-SVZ during a basal state in our model, as injury or another stimulus may be required for these cells to migrate to the SVZ.

In summary, we have identified a largely quiescent, multipotent stem cell population in the adult mouse brain which expresses TERT. These cells are localized to various regions of the adult mouse brain and give rise to numerous adult-born mature cell types in the basal state of tissue turnover and regeneration. Surprisingly, only a small population of TERT+ cells and their progeny were found in the traditional V-SVZ, RMS, and OB neurogenic niches, while no population of TERT+ cells existed in the SGZ. We hypothesize that the low number of GFP+ and mGFP+ cells identified in these classically neurogenic niches indicates that TERT+ cells may be a separate population of ASCs in the adult mammalian brain, or that they could give rise to the stem/progenitor cells that have been previously studied in adult neurogenesis in these niches, either earlier in development or in response to adult stimuli that we did not investigate in this particular study. It is also possible that these classical niches house activated stem cells or committed progenitors, but not ASCs, if TERT marks the sole ASC population in the brain. Intriguingly, TERT+ ASCs express only a subset of the markers previously used for ANSCs, which could be because they are a true multipotent stem cell and not a committed neural/glial progenitor cell.

## Acknowledgements

The authors wish to thank Bethany Dudley Miles, Jillian Gori, Ashley Ronzo, and Cameron Ford, for technical assistance, Tianyi Tao for a critical read of the manuscript, as well as Brenda Kennedy-Wade for management of the UMaine small animal research facility, Will Schott and the Jackson Laboratory FACS Core, the OSU Confocal Imaging Cores (funded by: P30 NS104177 & S10 OD026842, and NIH grant 1S10OD026842-01), and the OSU FACS Core (funded by: CCSG: P30CA016058). We thank Ashley Webb, Liz Kirby, Thane Fremouw, and Martin Pera for helpful discussions. KT, GJ and AB are the guarantors of the work in this manuscript

## Supplementary material

‘Supplementary material is available at *Brain* online’.

## Abbreviations

3V: Third Ventricle

4V: Fourth Ventricle

aASC: Activated Adult Stem Cell

Ara-C: Cytosine-beta-D-arabinofuranoside

ARC: Arcuate Nucleus

bFGF: Basic Fibroblast Growth Factor

BMP: Bone Morphogenetic Protein

BMPR1A: BMP Receptor 1A

ChP: Choroid Plexus

CSF: Cerebrospinal Fluid

D3V: Dorsal Third Ventricle

DCX: Doublecortin

DG: Dentate Gyrus

EGF: Epidermal Growth Factor

FACS: Fluorescence Activated Cell Sorting

FOXO: Forkhead box O

GCL: Granule Cell Layer

GFAP: Glial Fibrillary Acidic Protein

GLAST: Glutamate Aspartate Transporter

IoC: Islands of Calleja

ME: Median Eminence

MiL: Mitral Layer

NeuN: Neuronal Nuclei

NG2: Neuron-Glial Antigen 2

NPC: Neural Progenitor Cell

OB: Olfactory Bulb

Olig2: Oligodendrocyte Transcription Factor 2

ONL: Olfactory Nerve Layer

PGL: Periglomerular Layer

qANSC: (Quiescent) Adult Neural Stem Cell

qASC: (Quiescent) Adult Stem Cell

RMS: Rostral Migratory Stream

SGZ: Subgranular Zone

Sox2: SRY-Box Transcription Factor 2

TAC: Transit Amplifying Cell

TERT: Telomerase Reverse Transcriptase

V-SVZ: Ventricular-Subventricular Zone

## References

1. Ming GL, Song H. Adult neurogenesis in the mammalian brain: significant answers and significant questions. Neuron. May 26 2011;70(4):687–702. doi:10.1016/j.neuron.2011.05.001

2. Jurkowski MP, Bettio L, E KW, Patten A, Yau SY, Gil-Mohapel J. Beyond the Hippocampus and the SVZ: Adult Neurogenesis Throughout the Brain. Front Cell Neurosci. 2020;14:576444. doi:10.3389/fncel.2020.576444

3. Breault DT, Min IM, Carlone DL, et al. Generation of mTert-GFP mice as a model to identify and study tissue progenitor cells. Proc Natl Acad Sci U S A. Jul 29 2008;105(30):10420–5. doi:10.1073/pnas.0804800105

4. Carlone DL, Riba-Wolman RD, Deary LT, et al. Telomerase expression marks transitional growth-associated skeletal progenitor/stem cells. Stem Cells. Mar 2021;39(3):296–305. doi:10.1002/stem.3318

5. Cousins FL, O DF, Ong YR, Breault DT, Deane JA, Gargett CE. Telomerase Reverse Transcriptase Expression in Mouse Endometrium During Reepithelialization and Regeneration in a Menses-Like Model. Stem Cells Dev. Jan 1 2019;28(1):1–12. doi:10.1089/scd.2018.0133

6. Deane JA, Ong YR, Cain JE, et al. The mouse endometrium contains epithelial, endothelial and leucocyte populations expressing the stem cell marker telomerase reverse transcriptase. Mol Hum Reprod. Apr 2016;22(4):272–84. doi:10.1093/molehr/gav076

7. Lin S, Nascimento EM, Gajera CR, et al. Distributed hepatocytes expressing telomerase repopulate the liver in homeostasis and injury. Nature. Apr 2018;556(7700):244-248. doi:10.1038/s41586-018-0004-7

8. Lynes MD DC, KL Townsend, DT Braeault, Y-H Tseng. Telomerase reverse transcriptase expression marks a population of rare adipose tissue stem cells. 2021 (accepted manuscript);

9. Martin-Rivera L, Herrera E, Albar JP, Blasco MA. Expression of mouse telomerase catalytic subunit in embryos and adult tissues. Proc Natl Acad Sci U S A. Sep 1 1998;95(18):10471–6. doi:10.1073/pnas.95.18.10471

10. Montgomery RK, Carlone DL, Richmond CA, et al. Mouse telomerase reverse transcriptase (mTert) expression marks slowly cycling intestinal stem cells. Proc Natl Acad Sci U S A. Jan 4 2011;108(1):179–84. doi:10.1073/pnas.1013004108

11. Richardson GD, Breault D, Horrocks G, Cormack S, Hole N, Owens WA. Telomerase expression in the mammalian heart. FASEB J. Dec 2012;26(12):4832–40. doi:10.1096/fj.12-208843

12. Song J, Czerniak S, Wang T, et al. Characterization and fate of telomerase-expressing epithelia during kidney repair. J Am Soc Nephrol. Dec 2011;22(12):2256–65. doi:10.1681/ASN.2011050447

13. Wright WE, Piatyszek MA, Rainey WE, Byrd W, Shay JW. Telomerase activity in human germline and embryonic tissues and cells. Dev Genet. 1996;18(2):173–9. doi:10.1002/(SICI)1520-6408(1996)18:2<173::AID-DVG10>3.0.CO;2-3

14. Greenberg RA, Allsopp RC, Chin L, Morin GB, DePinho RA. Expression of mouse telomerase reverse transcriptase during development, differentiation and proliferation. Oncogene. Apr 2 1998;16(13):1723–30. doi:10.1038/sj.onc.1201933

15. Caporaso GL, Lim DA, Alvarez-Buylla A, Chao MV. Telomerase activity in the subventricular zone of adult mice. Mol Cell Neurosci. Aug 2003;23(4):693–702. doi:10.1016/s1044-7431(03)00103-9

16. Zhou QG, Hu Y, Wu DL, et al. Hippocampal telomerase is involved in the modulation of depressive behaviors. J Neurosci. Aug 24 2011;31(34):12258–69. doi:10.1523/JNEUROSCI.0805-11.2011

17. Kruk PA, Balajee AS, Rao KS, Bohr VA. Telomere reduction and telomerase inactivation during neuronal cell differentiation. Biochem Biophys Res Commun. Jul 16 1996;224(2):487–92. doi:10.1006/bbrc.1996.1054

18. Miura T, Katakura Y, Yamamoto K, et al. Neural stem cells lose telomerase activity upon differentiating into astrocytes. Cytotechnology. Jul 2001;36(1-3):137–44. doi:10.1023/A:1014016315003

19. Ferron SR, Marques-Torrejon MA, Mira H, et al. Telomere shortening in neural stem cells disrupts neuronal differentiation and neuritogenesis. J Neurosci. Nov 18 2009;29(46):14394–407. doi:10.1523/JNEUROSCI.3836-09.2009

20. Jaskelioff M, Muller FL, Paik JH, et al. Telomerase reactivation reverses tissue degeneration in aged telomerase-deficient mice. Nature. Jan 6 2011;469(7328):102–6. doi:10.1038/nature09603

21. Saha B, Peron S, Murray K, Jaber M, Gaillard A. Cortical lesion stimulates adult subventricular zone neural progenitor cell proliferation and migration to the site of injury. Stem Cell Res. Nov 2013;11(3):965–77. doi:10.1016/j.scr.2013.06.006

22. Usta SN, Scharer CD, Xu J, Frey TK, Nash RJ. Chemically defined serum-free and xeno-free media for multiple cell lineages. Ann Transl Med. Oct 2014;2(10):97. doi:10.3978/j.issn.2305-5839.2014.09.05

23. Renier N, Wu Z, Simon DJ, Yang J, Ariel P, Tessier-Lavigne M. iDISCO: a simple, rapid method to immunolabel large tissue samples for volume imaging. Cell. Nov 6 2014;159(4):896–910. doi:10.1016/j.cell.2014.10.010

24. Decimo I, Fumagalli G, Berton V, Krampera M, Bifari F. Meninges: from protective membrane to stem cell niche. Am J Stem Cells. 2012;1(2):92–105.

25. Itokazu Y, Kitada M, Dezawa M, et al. Choroid plexus ependymal cells host neural progenitor cells in the rat. Glia. Jan 1 2006;53(1):32–42. doi:10.1002/glia.20255

26. Jurga AM, Paleczna M, Kuter KZ. Overview of General and Discriminating Markers of Differential Microglia Phenotypes. Front Cell Neurosci. 2020;14:198. doi:10.3389/fncel.2020.00198

27. Reynolds BA, Rietze RL. Neural stem cells and neurospheres--re-evaluating the relationship. Nat Methods. May 2005;2(5):333–6. doi:10.1038/nmeth758

28. Zhang J, Jiao J. Molecular Biomarkers for Embryonic and Adult Neural Stem Cell and Neurogenesis. Biomed Res Int. 2015;2015:727542. doi:10.1155/2015/727542

29. Jensen GS, Leon-Palmer NE, Townsend KL. Bone morphogenetic proteins (BMPs) in the central regulation of energy balance and adult neural plasticity. Metabolism. Jul 29 2021;123:154837. doi:10.1016/j.metabol.2021.154837

30. Nishiyama A, Watanabe M, Yang Z, Bu J. Identity, distribution, and development of polydendrocytes: NG2-expressing glial cells. J Neurocytol. Jul-Aug 2002;31(6-7):437–55. doi:10.1023/a:1025783412651

31. Doetsch F, Caille I, Lim DA, Garcia-Verdugo JM, Alvarez-Buylla A. Subventricular zone astrocytes are neural stem cells in the adult mammalian brain. Cell. Jun 11 1999;97(6):703–16. doi:10.1016/s0092-8674(00)80783-7

32. Carlone DL. Identifying Adult Stem Cells Using Cre-Mediated Lineage Tracing. Curr Protoc Stem Cell Biol. Feb 3 2016;36:5a.2.1-5a.2.18. doi:10.1002/9780470151808.sc05a02s36

33. Muzumdar MD, Tasic B, Miyamichi K, Li L, Luo L. A global double-fluorescent Cre reporter mouse. Genesis. Sep 2007;45(9):593–605. doi:10.1002/dvg.20335

34. Kempermann G. Why new neurons? Possible functions for adult hippocampal neurogenesis. J Neurosci. Feb 1 2002;22(3):635–8.

35. Kokoeva MV, Yin H, Flier JS. Neurogenesis in the hypothalamus of adult mice: potential role in energy balance. Science. Oct 28 2005;310(5748):679–83. doi:10.1126/science.1115360

36. Lugert S, Basak O, Knuckles P, et al. Quiescent and active hippocampal neural stem cells with distinct morphologies respond selectively to physiological and pathological stimuli and aging. Cell Stem Cell. May 7 2010;6(5):445–56. doi:10.1016/j.stem.2010.03.017

37. Ortega-Perez I, Murray K, Lledo PM. The how and why of adult neurogenesis. J Mol Histol. Dec 2007;38(6):555–62. doi:10.1007/s10735-007-9114-5

38. Petreanu L, Alvarez-Buylla A. Maturation and death of adult-born olfactory bulb granule neurons: role of olfaction. J Neurosci. Jul 15 2002;22(14):6106–13. doi:20026588

39. Lledo PM, Valley M. Adult Olfactory Bulb Neurogenesis. Cold Spring Harb Perspect Biol. Aug 1 2016;8(8) doi:10.1101/cshperspect.a018945

40. Lee DA, Yoo S, Pak T, et al. Dietary and sex-specific factors regulate hypothalamic neurogenesis in young adult mice. Front Neurosci. 2014;8:157. doi:10.3389/fnins.2014.00157

41. Kokoeva MV, Yin H, Flier JS. Evidence for constitutive neural cell proliferation in the adult murine hypothalamus. J Comp Neurol. Nov 10 2007;505(2):209–20. doi:10.1002/cne.21492

42. Goodman T, Hajihosseini MK. Hypothalamic tanycytes-masters and servants of metabolic, neuroendocrine, and neurogenic functions. Front Neurosci. 2015;9:387. doi:10.3389/fnins.2015.00387

43. Ahlfeld J, Filser S, Schmidt F, et al. Neurogenesis from Sox2 expressing cells in the adult cerebellar cortex. Sci Rep. Jul 21 2017;7(1):6137. doi:10.1038/s41598-017-06150-x

44. Nakagomi T, Molnar Z, Nakano-Doi A, et al. Ischemia-induced neural stem/progenitor cells in the pia mater following cortical infarction. Stem Cells Dev. Dec 2011;20(12):2037–51. doi:10.1089/scd.2011.0279

45. Decimo I, Dolci S, Panuccio G, Riva M, Fumagalli G, Bifari F. Meninges: A Widespread Niche of Neural Progenitors for the Brain. Neuroscientist. Sep 16 2020:1073858420954826. doi:10.1177/1073858420954826

46. Tumani H, Huss A, Bachhuber F. The cerebrospinal fluid and barriers - anatomic and physiologic considerations. Handb Clin Neurol. 2017;146:21–32. doi:10.1016/B978-0-12-804279-3.00002-2

47. Silva-Vargas V, Maldonado-Soto AR, Mizrak D, Codega P, Doetsch F. Age-Dependent Niche Signals from the Choroid Plexus Regulate Adult Neural Stem Cells. Cell Stem Cell. Nov 3 2016;19(5):643–652. doi:10.1016/j.stem.2016.06.013

48. Lun MP, Monuki ES, Lehtinen MK. Development and functions of the choroid plexus-cerebrospinal fluid system. Nat Rev Neurosci. Aug 2015;16(8):445–57. doi:10.1038/nrn3921

49. Feinstein P, Bozza T, Rodriguez I, Vassalli A, Mombaerts P. Axon guidance of mouse olfactory sensory neurons by odorant receptors and the beta2 adrenergic receptor. Cell. Jun 11 2004;117(6):833–46. doi:10.1016/j.cell.2004.05.013

50. Schwob JE, Farber NB, Gottlieb DI. Neurons of the olfactory epithelium in adult rats contain vimentin. J Neurosci. Jan 1986;6(1):208–17.

51. Paton JA, Nottebohm FN. Neurons generated in the adult brain are recruited into functional circuits. Science. Sep 7 1984;225(4666):1046-8. doi:10.1126/science.6474166

52. Richards LJ, Kilpatrick TJ, Bartlett PF. De novo generation of neuronal cells from the adult mouse brain. Proc Natl Acad Sci U S A. Sep 15 1992;89(18):8591–5. doi:10.1073/pnas.89.18.8591

53. Reynolds BA, Weiss S. Generation of neurons and astrocytes from isolated cells of the adult mammalian central nervous system. Science. Mar 27 1992;255(5052):1707-10. doi:10.1126/science.1553558

54. Eriksson PS, Perfilieva E, Bjork-Eriksson T, et al. Neurogenesis in the adult human hippocampus. Nat Med. Nov 1998;4(11):1313–7. doi:10.1038/3305

55. Lim DA, Alvarez-Buylla A. The Adult Ventricular-Subventricular Zone (V-SVZ) and Olfactory Bulb (OB) Neurogenesis. Cold Spring Harb Perspect Biol. May 2 2016;8(5)doi:10.1101/cshperspect.a018820

56. Ernst A, Alkass K, Bernard S, et al. Neurogenesis in the striatum of the adult human brain. Cell. Feb 27 2014;156(5):1072–83. doi:10.1016/j.cell.2014.01.044

57. Abbott LC, Nigussie F. Adult neurogenesis in the mammalian dentate gyrus. Anat Histol Embryol. Jan 2020;49(1):3–16. doi:10.1111/ahe.12496

58. Petrik D, Encinas JM. Perspective: Of Mice and Men - How Widespread Is Adult Neurogenesis? Front Neurosci. 2019;13:923. doi:10.3389/fnins.2019.00923

59. Lin S, Nascimento EM, Gajera CR, et al. Distributed hepatocytes expressing telomerase repopulate the liver in homeostasis and injury. Nature. Apr 2018;556<otherinfo>(7700):244-248. doi:10.1038/s41586-018-0004-7</otherinfo>

60. Lynes MD, Carlone DL, Townsend KL, Breault DT, Tseng Y-H. Telomerase reverse transcriptase expression marks a population of rare adipose tissue stem cells. Stem Cells. 2021 (accepted manuscript);

61. Liu JP. Studies of the molecular mechanisms in the regulation of telomerase activity. FASEB J. Dec 1999;13(15):2091–104. doi:10.1096/fasebj.13.15.2091

62. Spilsbury A, Miwa S, Attems J, Saretzki G. The role of telomerase protein TERT in Alzheimer’s disease and in tau-related pathology in vitro. J Neurosci. Jan 28 2015;35(4):1659–74. doi:10.1523/JNEUROSCI.2925-14.2015

63. Ong YR, Cousins FL, Yang X, et al. Bone Marrow Stem Cells Do Not Contribute to Endometrial Cell Lineages in Chimeric Mouse Models. Stem Cells. Jan 2018;36(1):91–102. doi:10.1002/stem.2706

64. Zvereva MI, Shcherbakova DM, Dontsova OA. Telomerase: structure, functions, and activity regulation. Biochemistry (Mosc). Dec 2010;75(13):1563–83. doi:10.1134/s0006297910130055

65. Zheng Q, Huang J, Wang G. Mitochondria, Telomeres and Telomerase Subunits. Front Cell Dev Biol. 2019;7:274. doi:10.3389/fcell.2019.00274

66. Jose SS, Tidu F, Burilova P, Kepak T, Bendickova K, Fric J. The Telomerase Complex Directly Controls Hematopoietic Stem Cell Differentiation and Senescence in an Induced Pluripotent Stem Cell Model of Telomeropathy. Original Research. Frontiers in Genetics. 2018-August-29 2018;9doi:10.3389/fgene.2018.00345

67. Yin L, Hubbard AK, Giardina C. NF-kappa B regulates transcription of the mouse telomerase catalytic subunit. J Biol Chem. Nov 24 2000;275(47):36671–5. doi:10.1074/jbc.M007378200

68. Charvet CJ, Finlay BL. Comparing Adult Hippocampal Neurogenesis Across Species: Translating Time to Predict the Tempo in Humans. Front Neurosci. 2018;12:706. doi:10.3389/fnins.2018.00706

69. Cipriani S, Ferrer I, Aronica E, et al. Hippocampal Radial Glial Subtypes and Their Neurogenic Potential in Human Fetuses and Healthy and Alzheimer’s Disease Adults. Cereb Cortex. Jul 1 2018;28(7):2458–2478. doi:10.1093/cercor/bhy096

70. Boldrini M, Fulmore CA, Tartt AN, et al. Human Hippocampal Neurogenesis Persists throughout Aging. Cell Stem Cell. Apr 5 2018;22(4):589–599 e5. doi:10.1016/j.stem.2018.03.015

71. Knoth R, Singec I, Ditter M, et al. Murine features of neurogenesis in the human hippocampus across the lifespan from 0 to 100 years. PLoS One. Jan 29 2010;5(1):e8809. doi:10.1371/journal.pone.0008809

72. Moreno-Jimenez EP, Terreros-Roncal J, Flor-Garcia M, Rabano A, Llorens-Martin M. Evidences for Adult Hippocampal Neurogenesis in Humans. J Neurosci. Mar 24 2021;41(12):2541–2553. doi:10.1523/JNEUROSCI.0675-20.2020

73. Dennis CV, Suh LS, Rodriguez ML, Kril JJ, Sutherland GT. Human adult neurogenesis across the ages: An immunohistochemical study. Neuropathol Appl Neurobiol. Dec 2016;42(7):621–638. doi:10.1111/nan.12337

74. Sorrells SF, Paredes MF, Cebrian-Silla A, et al. Human hippocampal neurogenesis drops sharply in children to undetectable levels in adults. Nature. Mar 15 2018;555(7696):377-381. doi:10.1038/nature25975

75. Paredes MF, Sorrells SF, Cebrian-Silla A, et al. Does Adult Neurogenesis Persist in the Human Hippocampus? Cell Stem Cell. Dec 6 2018;23(6):780–781. doi:10.1016/j.stem.2018.11.006

76. Sorrells SF, Paredes MF, Zhang Z, et al. Positive Controls in Adults and Children Support That Very Few, If Any, New Neurons Are Born in the Adult Human Hippocampus. J Neurosci. Mar 24 2021;41(12):2554–2565. doi:10.1523/JNEUROSCI.0676-20.2020

77. Flor-Garcia M, Terreros-Roncal J, Moreno-Jimenez EP, Avila J, Rabano A, Llorens-Martin M. Unraveling human adult hippocampal neurogenesis. Nat Protoc. Feb 2020;15(2):668–693. doi:10.1038/s41596-019-0267-y

78. Gandhi S, Gupta J, Tripathi PP. The Curious Case of Human Hippocampal Neurogenesis. ACS Chem Neurosci. Mar 20 2019;10(3):1131–1132. doi:10.1021/acschemneuro.9b00063

79. Lucassen PJ, Fitzsimons CP, Salta E, Maletic-Savatic M. Adult neurogenesis, human after all (again): Classic, optimized, and future approaches. Behav Brain Res. Mar 2 2020;381:112458. doi:10.1016/j.bbr.2019.112458

80. Seki T. Understanding the Real State of Human Adult Hippocampal Neurogenesis From Studies of Rodents and Non-human Primates. Front Neurosci. 2020;14:839. doi:10.3389/fnins.2020.00839

81. Kempermann G, Gage FH, Aigner L, et al. Human Adult Neurogenesis: Evidence and Remaining Questions. Cell Stem Cell. Jul 5 2018;23(1):25–30. doi:10.1016/j.stem.2018.04.004

82. Klapper W, Shin T, Mattson MP. Differential regulation of telomerase activity and TERT expression during brain development in mice. J Neurosci Res. May 1 2001;64(3):252–60. doi:10.1002/jnr.1073

83. Horikawa I, Chiang YJ, Patterson T, et al. Differential cis-regulation of human versus mouse TERT gene expression in vivo: identification of a human-specific repressive element. Proc Natl Acad Sci U S A. Dec 20 2005;102(51):18437–42. doi:10.1073/pnas.0508964102

84. Ishaq A, Hanson PS, Morris CM, Saretzki G. Telomerase Activity is Downregulated Early During Human Brain Development. Genes (Basel). Jun 16 2016;7(6)doi:10.3390/genes7060027

85. Ulaner GA, Giudice LC. Developmental regulation of telomerase activity in human fetal tissues during gestation. Mol Hum Reprod. Sep 1997;3(9):769–73. doi:10.1093/molehr/3.9.769

86. Zhou QG, Liu MY, Lee HW, et al. Hippocampal TERT Regulates Spatial Memory Formation through Modulation of Neural Development. Stem Cell Reports. Aug 8 2017;9(2):543–556. doi:10.1016/j.stemcr.2017.06.014

87. Cheung TH, Rando TA. Molecular regulation of stem cell quiescence. Nat Rev Mol Cell Biol. Jun 2013;14(6):329–40. doi:10.1038/nrm3591

88. Terzi MY, Izmirli M, Gogebakan B. The cell fate: senescence or quiescence. Mol Biol Rep. Nov 2016;43(11):1213–1220. doi:10.1007/s11033-016-4065-0

89. Fawal MA, Davy A. Impact of Metabolic Pathways and Epigenetics on Neural Stem Cells. Epigenet Insights. 2018;11:2516865718820946. doi:10.1177/2516865718820946

90. Liu MY, Nemes A, Zhou QG. The Emerging Roles for Telomerase in the Central Nervous System. Front Mol Neurosci. 2018;11:160. doi:10.3389/fnmol.2018.00160

