## Supplemental Figures for "Telomerase reverse transcriptase (TERT)-expressing cells mark a novel stem cell population in the adult mouse brain"

Suppl. Fig. S1

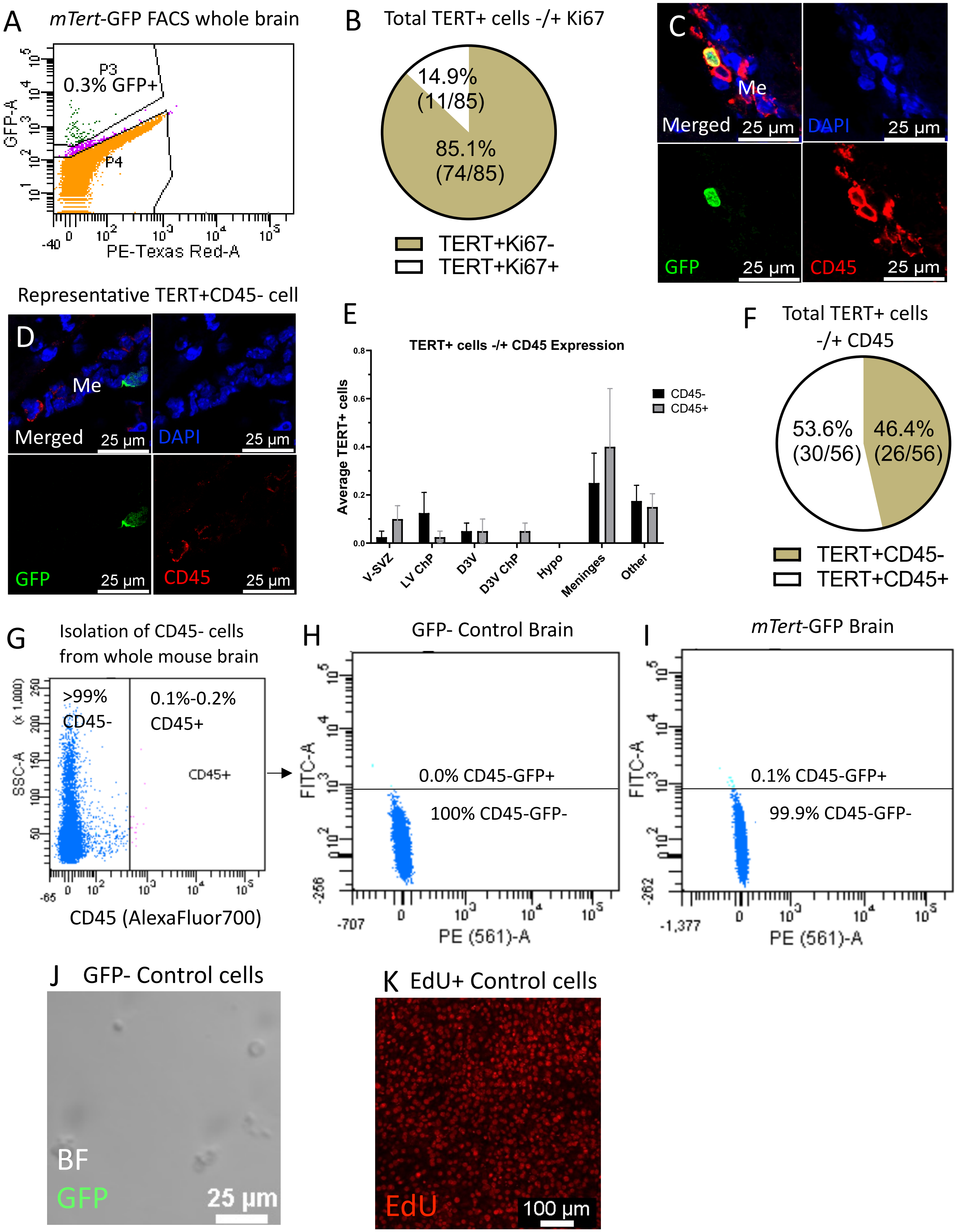

Suppl. Fig. S2

#### TERT+ Cell Co-Expression With Quiescent and Activated Stem Cell Markers

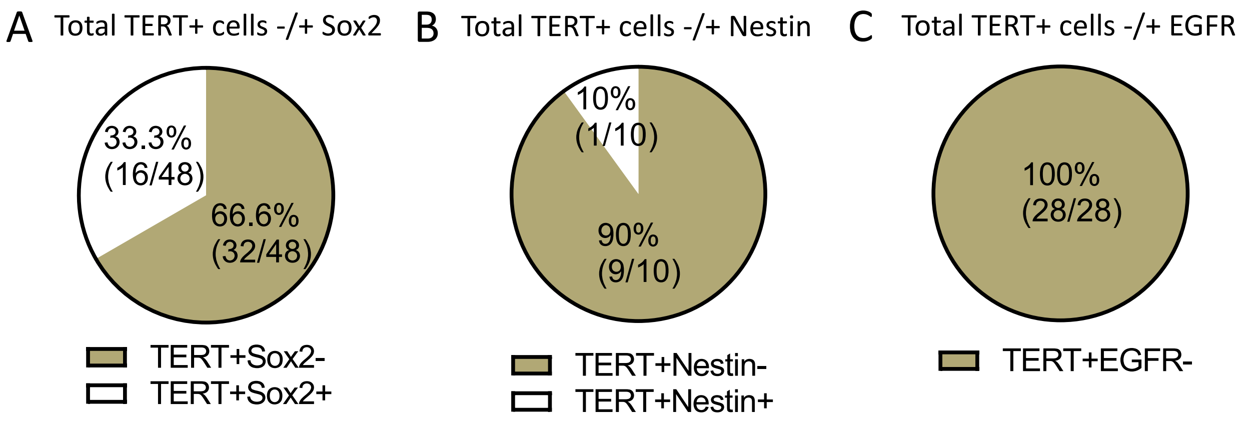


#### TERT+ Cell Co-expression with Neuroblast and Mature Neuron Markers

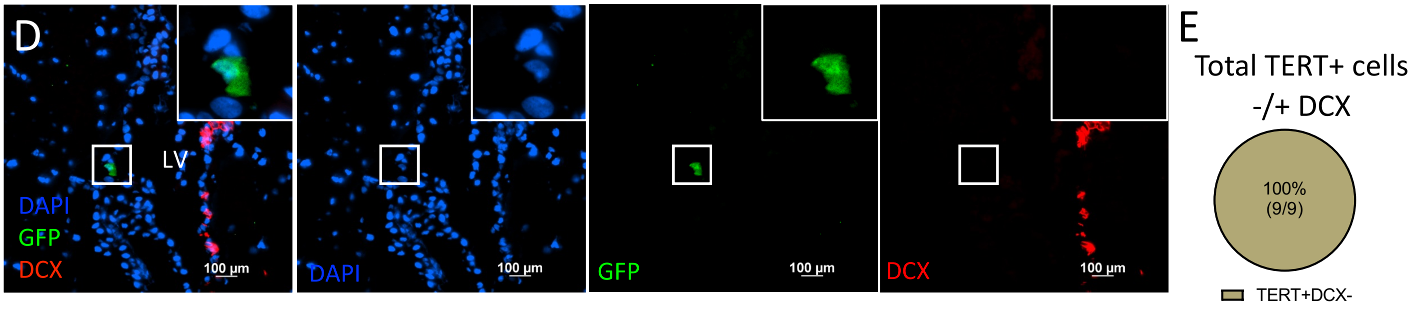


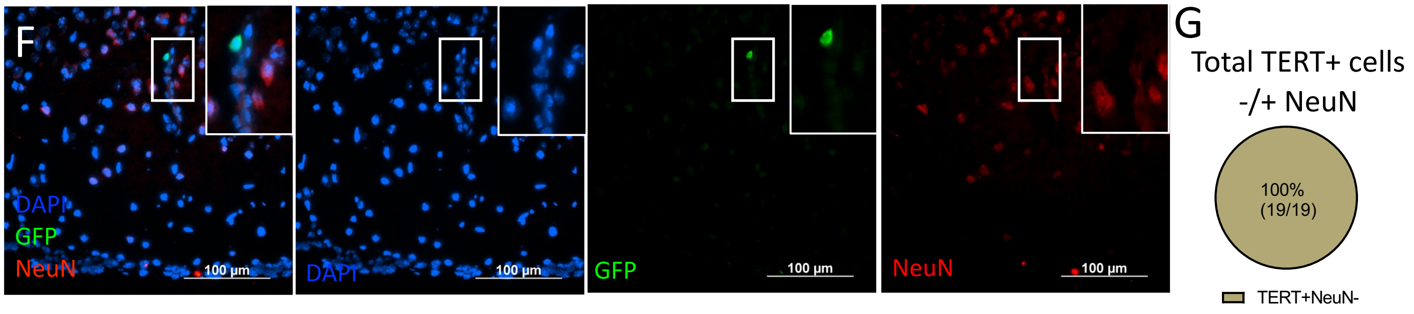


#### TERT+ Cell Co-expression with Bone Morphogenetic Protein Receptors

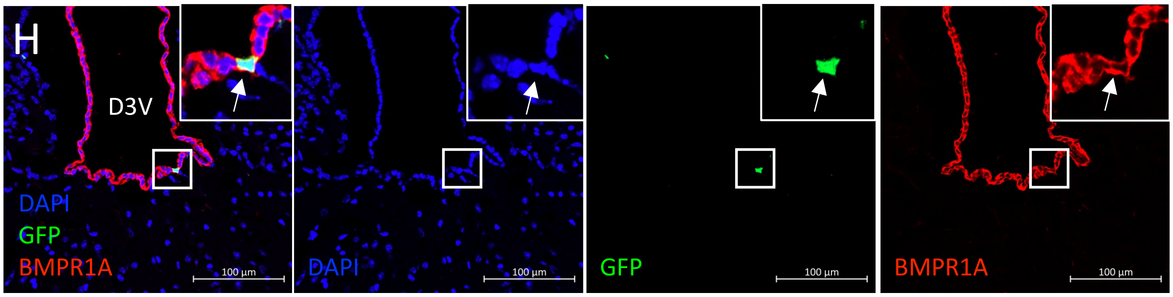

Suppl. Fig. S3

**TERT+ cells do not express markers of glial-committed progenitors**

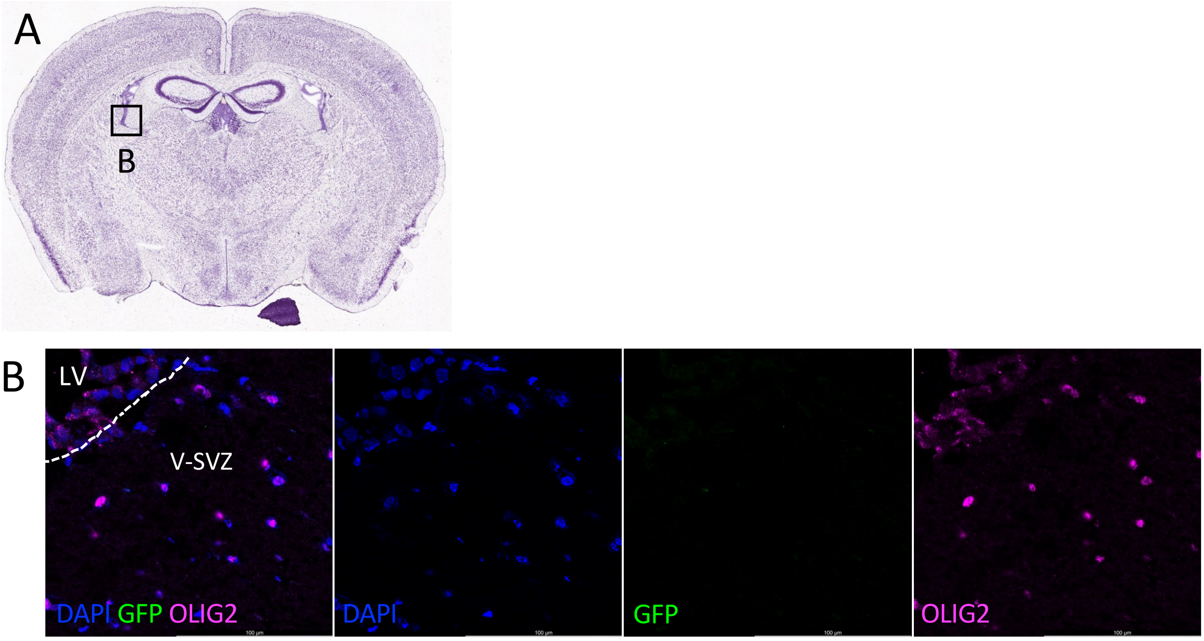

Suppl. Fig. S4

### TERT+ cells give rise to a heterogeneous population of cells in multiple plastic regions of the brain

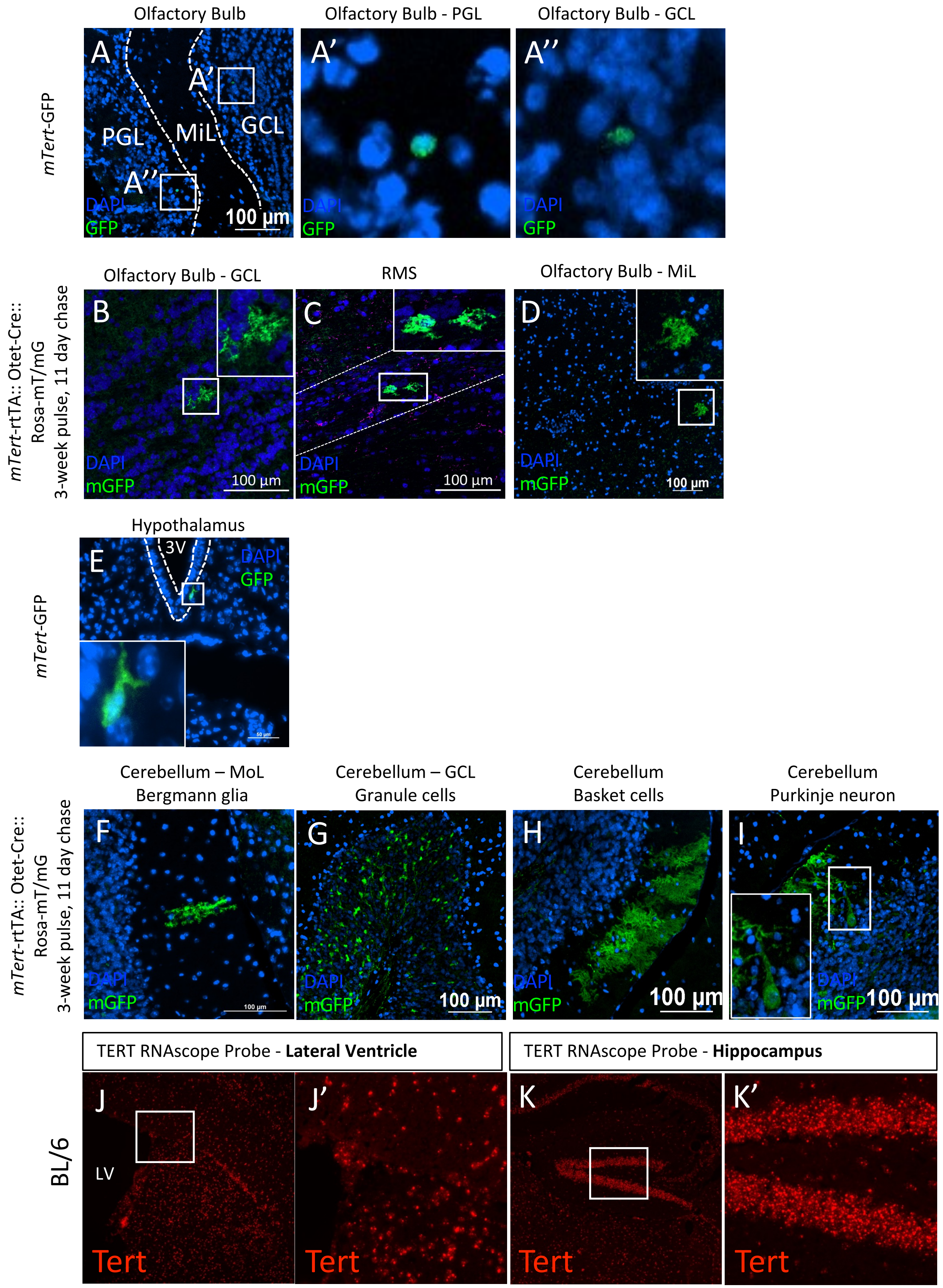

Supp. Fig. S5

### TERT+ cells give rise to a heterogeneous cell population in plastic regions of the brain

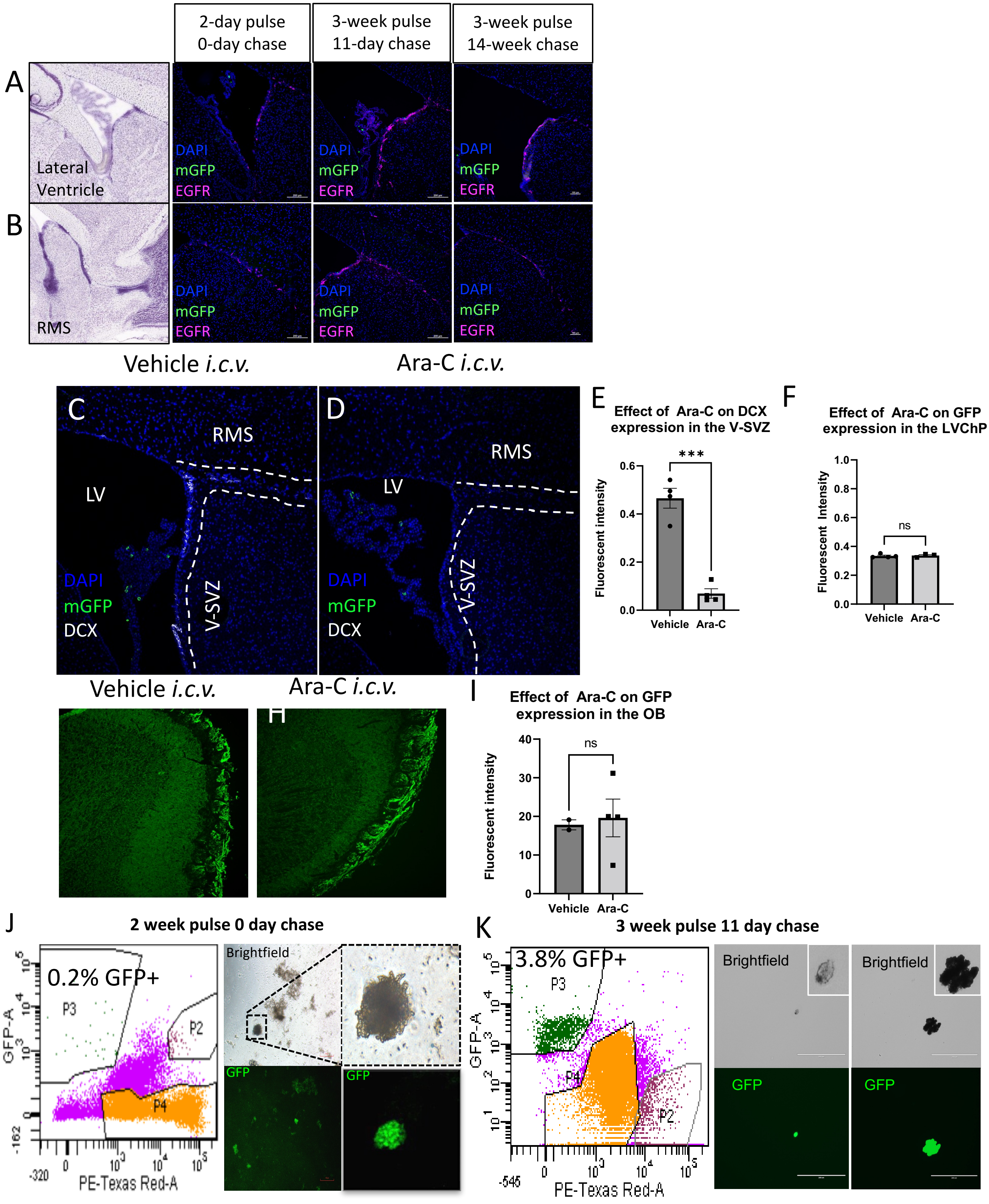

Suppl. Fig. S6

#### Comparison of Metabolic Activation of Lineage Tracing mTERT+ cells

5-day pulse, 0-day chase

##### 5-day *in Vitro*

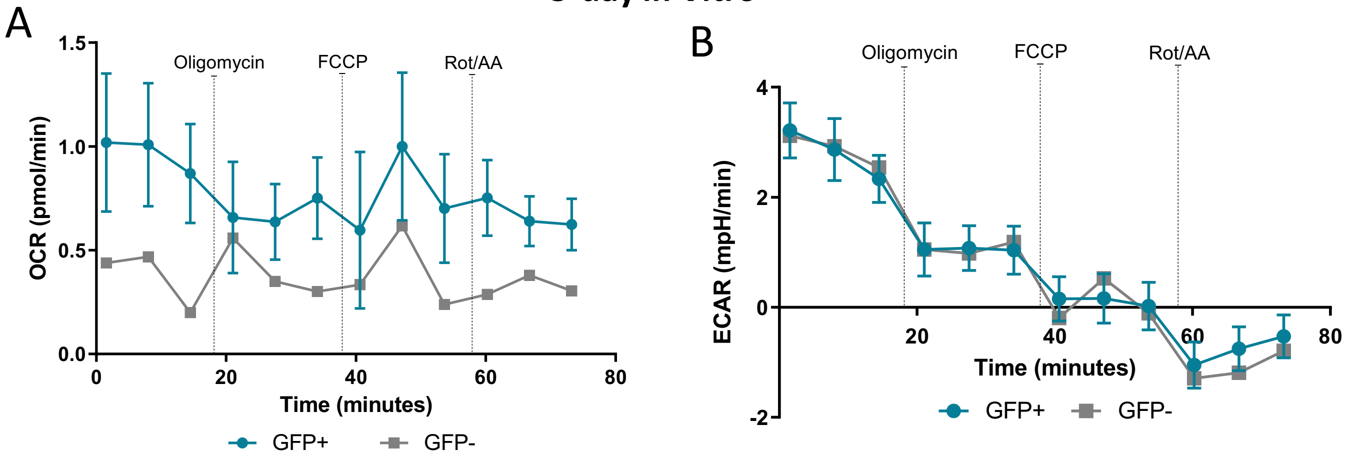


##### 15-day *In Vitro*- Neurospheres

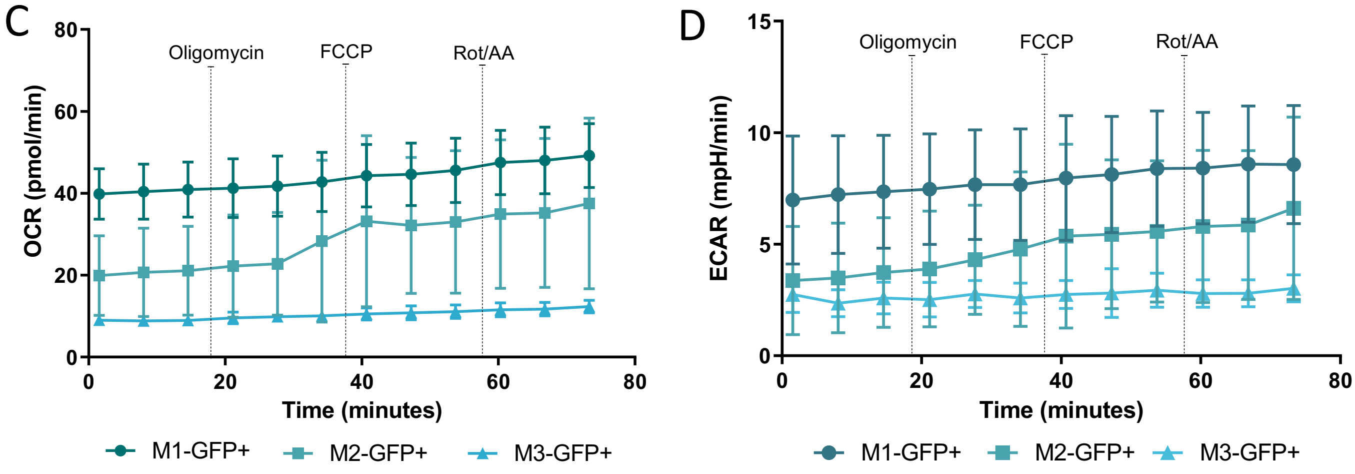

Suppl. Fig. S7

### TERT+ cells and neuronal cell types in the adult mouse brain

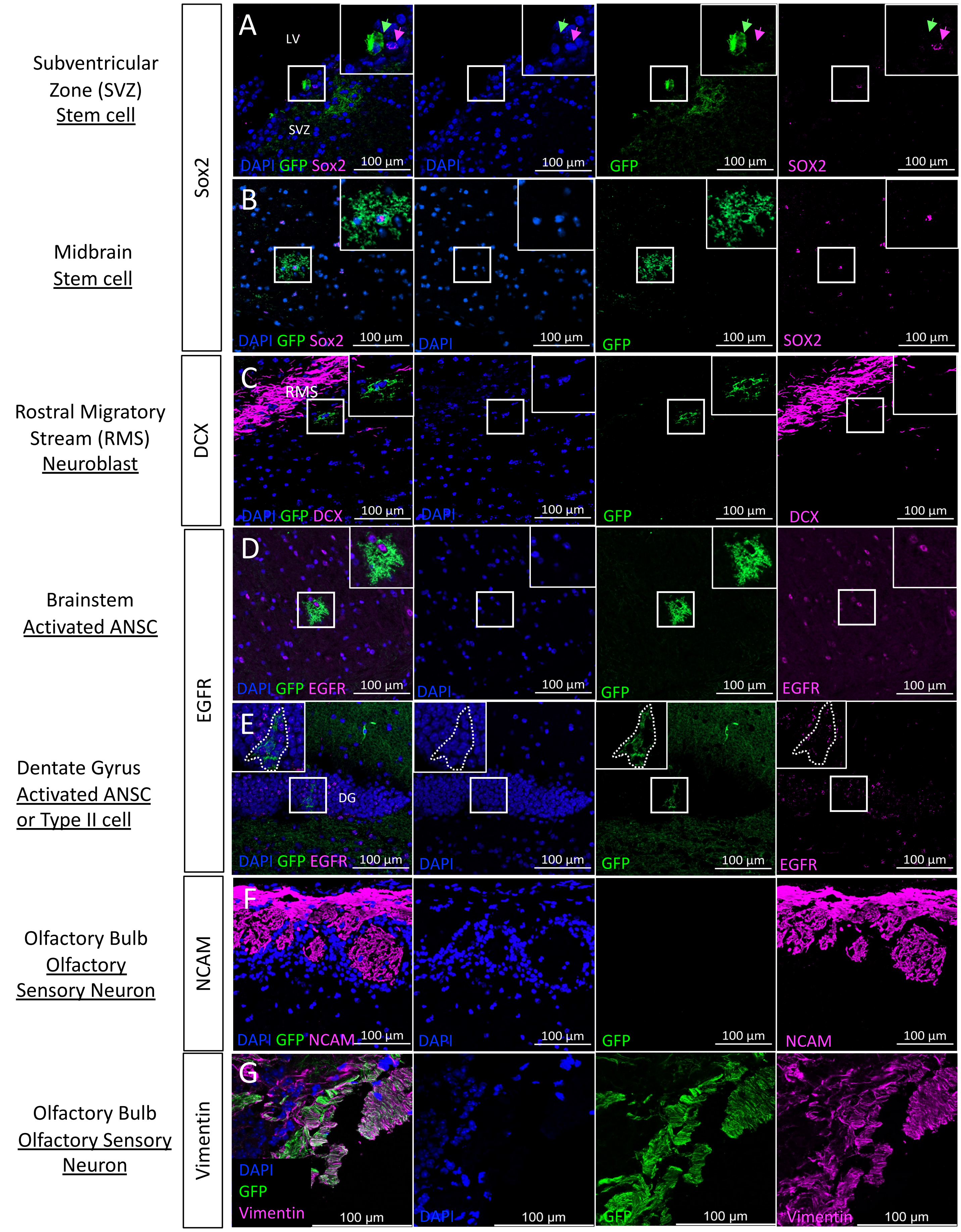

Suppl. Fig. S8

#### TERT+ cells and glial cell types in the adult mouse brain

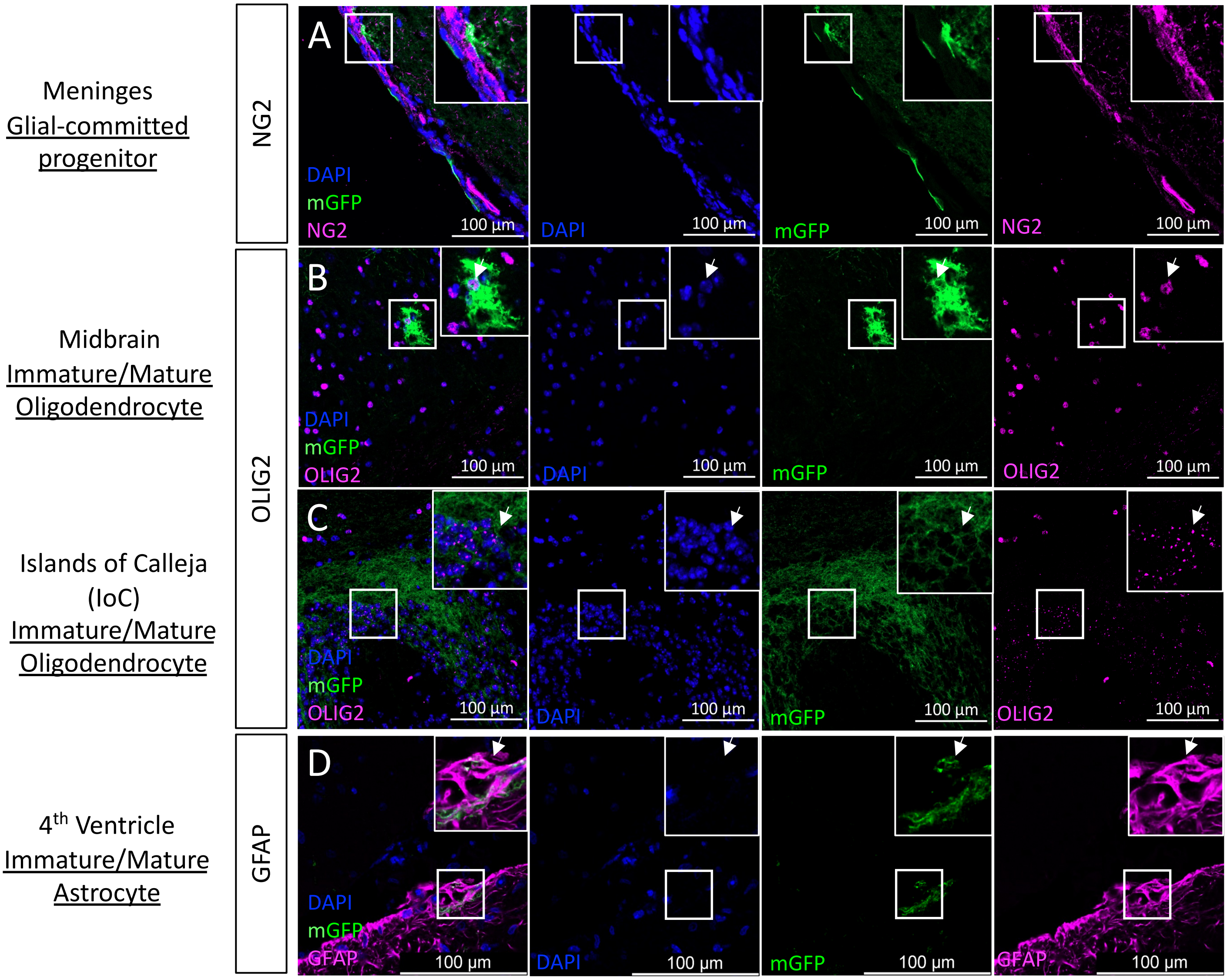
