## Supplemental Materials for "Telomerase reverse transcriptase (TERT)-expressing cells mark a novel stem cell population in the adult mouse brain"

### Supplementary Materials

#### Supplementary Figure Legends

**Supplementary Figure 1: TERT+ cells in the adult mouse brain are rare, mostly quiescent, and include a subpopulation of CD45+ immune cells. (A)** Representative fluorescent activated cells sorting (FACS) plot of GFP+ cells from sorted adult *mTert*-GFP full mouse brains ($N$ = 16 males, 9 females). P3: GFP+, P4: GFP-. PE-Texas Red-A used to gate for autofluorescence. **(B)** Quantification of total proliferative TERT+ cells across all tissues in the neurogenic enriched brain area stained for Ki67 from 10 *mTert-GFP* animals. ($N$ = 5 males, 5 females, $n$ = 8 sections per brain). **(C-D)** Representative images of TERT+CD45+ (C) and TERT+CD45- (D) cells identified via immunofluorescence in the adult *mTert*-GFP mouse brain (identified in $N$ = 5 of 5 males, 5 of 5 females, $n$ = 8 brain sections per mouse). **(E)** Quantification of co-staining for TERT-GFP and CD45 in *mTert*-GFP mouse brains. Cell quantification is graphed as the average number of cells per brain region across $n$ = 8 brain sections for each of $N$ = 5 males and 5 females. **(F)** Quantification of total proliferative TERT+ cells across all tissues in the neurogenic enriched brain area stained for Ki67 from 10 *mTert-GFP* animals. ($N$=5 males, 5 females, $n$ = 8 sections per brain). **(G-I)** Representative fluorescent activated cell sorting (FACS) plots showing identification of CD45- cells (G) followed by isolation of TERT-GFP+ cells from GFP- control brains (H) or full mTert-GFP full mouse brains (I) ($N$ = 6 males). **(J)** FACS sorted GFP- cells from *mTert*-GFP animals do not express GFP ($N$ = 2 males). **(K)** 3T3-F4421A cells treated with EdU for 24hr, used as a positive control of proliferation. **(L)** Table detailing the quantification of GFP+EdU+ cells across the total number of GFP+ cells per animal. ($N$ = 4 mice, $n$ = 1 well per brain).
Scale bars are 25μm.

**Supplementary Figure 2. TERT+ cells express markers of quiescent stem cells, but not of activated stem cells, neuronal precursors, or mature neurons. (A-C)** Quantification of total TERT+ cells across all tissues in the neurogenic enriched brain area stained for Sox2 (A), Nestin (B), or EGFR (C) from 10 *mTert-GFP* animals. ($N$ = 4-8 males, 6-9 females, $n$ = 4-8 sections per brain). **(D-G)** Representative images of TERT+ cells within the brain co-stained with DCX (D) with quantification across total TERT+ cells (E), or NeuN (F) with quantification across total TERT+ cells (G; for each marker, $N$ = 5 males, 5 females, $n$ = 4 sections per brain assessed). **(F)**

Brain section containing TERT+BMPR1A+ cell along D3V. 1 TERT+BMPR1A+ cell was identified across *N* = 5 males, 5 females, *n* = 8 sections per brain assessed.
Scale bars are all 100μm. Insets show indicated area at 2x digital zoom. LV: lateral ventricle, D3V: dorsal third ventricle.

**Supplementary Figure 3. TERT+ cells do not express markers of glial-committed progenitors or mature glia. (A)** Schematic of a coronal mouse brain indicating area of Fig. 3B (Allen Brain Atlas). **(B)** Positive control region for OLIG2 expression in the adult mouse brain. Scale bar is 100μm. LV: lateral ventricle, V-SVZ: ventricular-subventricular zone.

**Supplementary Figure 4. Lineage tracing reveals that TERT+ cells give rise to a heterogeneous population of cells in multiple plastic regions of the brain. (A)** Representative image of TERT+ cells in the OB of *mTert*-GFP animals, specifically the GCL (A') and PGL (A'') of mTert-GFP brains (identified in *N* = 2 of 5 mice [*N* = 2 of 3 males, 0 of 3 females], *n* = 8-14 sections per brain assessed). **(B)** Representative image of mGFP+ cell within the GCL of *mTert*-mTmG mice after a 3-week, 11-day lineage tracing experiment (identified in *N* = 8 of 10 mice [*N* = 3 of 4 males, 5 of 6 females], *n* = 13-23 sections per brain assessed). **(C)** Representative image of mGFP+ cell within the RMS of *mTert*-mTmG mice after a 3-week, 11-day lineage tracing experiment (identified in *N* = 3 of 10 mice [*N* = 2 of 4 males, 1 of 6 females], *n* = 1-5 sections per brain assessed). **(D)** mGFP+ cells within the MiL after lineage trace (identified in *N* = 2 of 10 mice [*N* = 1 of 4 males, 1 of 6 females], *n* = 13-22 sections per brain assessed. **(E)** Hypothalamic TERT+ cell within the tanycytic layers of the third ventricle (3V) (identified in *N*=1 of 11 mice [*N*= 1of 5 males, 0 of 6 females], *n* = 4-20 sections per brain assessed). **(F-I)** Representative images of mGFP+ cells following a 3-week, 11-day pulse-chase in the cerebellum identifying Bergmann glia (F), granule cells G), basket cells (H), and Purkinje neurons (I) (seen in *N* = 8-11 mice [N=4 of 4 males, 4-7 of 7 females], *n* = 4-19 sections per brain assessed).
Scale bars are 100μm.

**Supplementary Figure 5: Lineage tracing of TERT+ cells revealed low numbers of traced cells in certain adult brain niches, which were unaffected by mitotic inhibition. Lineage traced cells were able to form neurospheres. (A-B)** Representative images of each of the V-SVZ

(A), and RMS (B). Representative images of sagittal-view brain regions of interest were obtained from the Allen Brain Atlas. mTert-mTmG mouse brains were immunostained following either a: 2-day pulse, 0-day chase (*N* = 5 males, 5 females), 3-week pulse, 11-day chase (*N* = 4 males, 8 females), or 3-week pulse, 14-week chase (*N* = 6 females). **(C-D)** Representative image of the LV of a vehicle treated mouse (C) and Ara-C treated mouse (D). For both, *N* = 4 mice, *n* = 1 section per brain assessed. Dotted lines indicate areas of the V-SVZ and RMS where DCX expression is observed under basal conditions. **(E)** Fluorescent intensity of DCX signal in the V-SVZ of Ara-C or vehicle treated mice (*N* = 4 mice, *n* = 1 section per brain assessed). **(F)** Fluorescent intensity of GFP signal in the LV of Ara-C or vehicle treated mice quantified by frame of vision (*N* = 4 mice, *n* = 1 section per brain assessed). **(G-H)** Representative image of the OB of a vehicle treated mouse (G) and Ara-C treated mouse (H). For both, *N* = 4 mice, *n* = 1 section per brain assessed. **(I)** Fluorescent intensity of GFP signal in the OB of Ara-C or vehicle treated mice quantified by frame of vision (*N* = 4 mice, *n* = 1 section per brain assessed). **(J-K)** FACS analysis of 2-week pulse, 0-day chase (J; *N* = 1) and 3-week pulse, 11-day chase (K; *N* = 4) lineage-traced mouse brains and the ability of GFP+ cells to form neurospheres in culture. P3: GFP+, P2, P4: GFP-. PE-Texas Red-A used to analyze membrane tomato signal.

**Supplementary Figure 6: Mitochondrial respiration of lineage tracing of TERT+ cells after 5-day pulse followed by 0-day chase, revealed metabolic activity in traced cells after neurospheres formation. (A-B)** (A) OCR and (B) ECAR rates were measured of mTERT+ cells were separated through magnetic sorting following 5-day cell proliferation into a XFp cell culture miniplate. (n=5 animals, one per well) **(C-D)** (C) OCR and (D) ECAR rates were measured of mTERT+ cells following 15-day in cell culture, allowing these to form neurospheres which were dissociated prior the assay (n=03 animals, in duplicate). Oligomycin was added at 1.5 μM followed by FCCP at 1.5 μM and Rotenone/antimycin A at 0.5 μM.

**Supplementary Figure 7. Lineage trace of TERT cells leads to labeling of immature and mature neuronal cell types throughout the adult mouse brain. (A-B)** Representative images of 3-week, 11-day pulse-chased mTert-mTmG mouse brains stained for the stem cell marker Sox2 in the V-SVZ (A; *N* = 2 of 2 females, *n* = 2 sections per brain assessed) and midbrain (B; *N* = 2 of 2 males, 2 of 2 females, *n* = 2 sections per brain assessed). **(C)** Representative image of mGFP+DCX- cell in proximity to RMS after 3-week, 11-day pulse-chase (cells identified in *N* = 2 of 4 males, *n* = 1-5 sections per brain assessed). **(D)** Co-immunostaining with EGFR after 3-week, 11-day pulse-chase in the brainstem (*N* = 1 of 2 males, *n* = 2 sections per brain assessed). **(E)** TERT+EGFR cell identified within the DG after a 3-week, 14-week pulse-chase. 1 mGFP+ cell was found within the DG across 78 brain sections in 3-week, 14-week pulse-chased animals (*N* = 1 of 6 females, *n* = 13 sections per brain analyzed). No mGFP+ cells were identified in the DG in 2-day, 0-day or 3-week, 11-day pulse-chased animals (*N* = 4 males, 6-8 females, *n* = 2 sections per brain analyzed). **(F)** Representative image from a subset of female mice that showed little to no signal within the PGL or NL layers of the OB co-stained with the olfactory sensory neuron marker NCAM (*N* = 3 of 10 mice [*N* = 0 of 4 males, 3 of 6 females], *n* = 2 sections per brain assessed). **(G)** mGFP signal in the OB NL and PGL co-stained with Vimentin (*N* = 7 of 10 mice [*N* = 4 of 4 males, 3 of 6 females], *n* = 2 sections per brain assessed). All images taken in sagittal plane. Scale bars are 100µm. Insets show 2x digital magnification. Arrows indicate expression of GFP (green), other indicated marker (magenta), or co-staining (white).

**Supplementary Figure 8. Lineage trace of TERT cells leads to labeling of immature and mature glial cell types throughout the adult mouse brain. (A)** Representative image of co-immunostaining with NG2 in 3-week, 11-day pulse-chased mTert-mTmG lineage tracing mice (*N* = 5 of 5 mice [*N* = 2 of 2 males, 3 of 3 females], *n* = 2 sections per brain assessed). **(B-C)** Representative OLIG2 co-staining with GFP in 3-week, 11-day pulse-chased mTert-mTmG mouse brains in the midbrain (B; *N* = 3 of 4 mice [*N* = 2 of 2 males, 1 of 2 females], *n* = 2 sections per brain assessed) and IoC (C; *N* = 4 of 4 mice [*N* = 2 of 2 males, 2 of 2 females], *n* = 2 sections per brain assessed). **(D)** Representative co-staining of the astrocytic and ependymal marker GFAP in the 4V of 3-week, 11-day pulse-chased mTert-mTmG brains (identified in *N* = 2 of 10 mice [*N* = 2 of 4 males, 0 of 6 females], *n* = 2 sections per brain assessed). Scale bars are 100µm unless indicated otherwise. Insets show 2x digital zoom of indicated area. White arrows identify co-staining.

Supplementary Table 1: Genotyping Primers (5’ -> 3’)

| Gene | Primer 1 | Primer 2 | Primer 3 | Primer 4 |
|---|---|---|---|---|
| *mTert*-GFP | F: CAC ATG AAG CAG CAC GAC TT | R: AGT TCA CCT TGA TGC CGT TC | | |
| Cre | Cre F: CGTATATCCTGGCAGCGA | Cre R: CCGTTTGCCGGTCGTGGG | IRS-1 F: ACAGCGTGAATTTTGGAGTCAGAA | IRS-2 R: GTCTTGCTCAGCCTCGCTAT |
| *mTert*-rtTA | GCA CAG CAT TGC GGA CAT GC | CCC TCC ATG TGT GAC CAA GG | GCA GAA GCG CGG CCG TCT GG | |
| Rosa-mT/mG | WT-F: CTC TGC TGC CTC CTG GCT TCT | WT-R: CGA GGC GGA TCA CAA GCA AT' | Mut-R: TCA ATG GGC GGG GGT CGT T | |

**Key Resources Table**

| Reagent or Resource | Source | Identifier |
|---|---|---|
| **Antibodies** | | |
| goat anti-GFP | Abcam | ab6662, RRID:AB_305635 |
| Rabbit anti-GFP | Abcam | ab6556, RRID:AB_305564 |
| Rabbit anti-Ki67 | Abcam | ab15580, RRID:AB_443209 |
| Rat anti-CD45 (IF) | BD Bioscience | 550539, RRID:AB_2174426 |
| CD45 (FACS) | BioLegend | 103128, RRID:AB_493715 |
| Rabbit anti-BMPR1A | Invitrogen | 38-6000, RRID:AB_2533377 |
| Mouse anti-Nestin | Millipore-Sigma | MAB353, RRID:AB_94911 |
| Rabbit anti-Vimentin | Abcam | ab92547, RRID:AB_10562134 |
| Rabbit anti-EGFR | Millipore-Sigma | 06-847, RRID:AB_2096607 |
| Rabbit anti-GFAP | Cell Signaling Technologies | 12389, RRID:AB_2631098 |
| Rabbit anti-DCX | Abcam | ab18723, RRID:AB_732011 |
| Rabbit anti-NeuN | Abcam | ab177487, RRID:AB_2532109 |
| Rabbit anti-AQP1 | Abcam | ab15080, RRID:AB_2056839 |
| Rabbit anti-OLIG2 | Milipore-Sigma | AB9610, RRID:AB_570666 |
| Rabbit anti-NG2 | Milipore-Sigma | AB5320, RRID:AB_11213678 |
| Rabbit anti-Sox2 | Milipore-Sigma | AB5603, RRID:AB_2286686 |
| Rabbit anti-NCAM | Milipore-Sigma | AB5032, RRID:AB_2291692 |
| Rabbit anti-TUJ11 | Abcam | ab18207, RRID:AB_444319 |
| Mouse anti-PSA-NCAM, APC | Miltenyi Biotec | 130-120-43, RRID: AB 2727931 |
| CD45 MicroBeads, mouse | Miltenyi Biotec | 130-052-301 |
| **Biological Samples** | | |
| Adult brains, whole brain single cell suspensions, and cells grown from whole brain single cell suspensions from | This paper | N/A |

| | | |
|---|---|---|
| *mTert*-GFP, *mTert*-rtTA:: otet-Cre:: R26R(mT/mG), and *mTert*-rtTA:: otet-Cre:: Rosa-Luciferase mice | | |
| **Chemicals, Solutions, Peptides, and Recombinant Proteins** | | |
| Doxycycline Hyclate | Sigma-Aldrich | D9891 |
| Pronase | Millipore Sigma | 10165921001 |
| Sucrose | Millipore Sigma | S7903 |
| Artificial Cerebrospinal Fluid | Ecocyte | LRE-S-LSG-1000-1 |
| Fetal Bovine Serum | Gibco | A38401 |
| TRIzol Reagent | Ambion | 15596026 |
| Debris Removal Solution | Miltenyi | 130-109-398 |
| NeuroCult Basal Medium (Mouse and Rat) | Stem Cell Tech | 05700 |
| NeuroCult Proliferation Supplement (Mouse and Rat) | Stem Cell Tech | 05701 |
| Heparin Solution | Stem Cell Tech | 07980 |
| Cytosine-β-D-arabinoside | Millipore Sigma | C1768 |
| Human Recombinant bFGF | Stem Cell Tech | 78003 |
| Human Recombinant EGF | Stem Cell Tech | 78006 |
| Histochoice Molecular Biology Tissue Fixative | Amresco | H120 |
| Sudan (Typogen) Black | Millipore Sigma | 199664 |
| Triton X-100 | Bio-Rad | 1610407 |
| Tween-20 | Millipore Sigma | 655205 |
| Universal SSA SYBR Green | Bio-Rad | 1725271 |
| Optimal Cutting Temperature Solution | Sakura | 4583 |

| | | |
|---|---|---|
| Antigen Retrieval Solution | Agilent | S2367 |
| Blocking Solution | Millipore Sigma | 20773 |
| IHC Select TBS Rinse Buffer | Millipore Sigma | 20845 |
| Kwik-Sil Low Toxicity Silicone Adhesive | World Precision Instruments | KWIK-SIL |
| Heparin Sodium Salt from Porcine Mucosa | Sigma-Aldrich | H3393 |
| 10X PBS Solution | Teknova | P0496 |
| Dimethyl Sulfoxide | Sigma-Aldrich | D8418 |
| Dichloromethane | Sigma-Aldrich | 270997 |
| Donkey Serum | Sigma-Aldrich | D9663 |
| Tetrahydrofuran | Sigma-Aldrich | 186562 |
| Hydrogen Peroxide Solution | Sigma-Aldrich | 516813 |
| Paraformaldehyde | Sigma-Aldrich | P6148 |
| Glycine | Bio-Rad | 161-0718 |
| **Critical Commercial Assays** | | |
| Zymo Direct-zol RNA Microprep kit | Zymo | R2061 |
| High-Capacity cDNA Reverse Transcription Kit | Thermofisher | 4368813 |
| Adult Brain Dissociation Kit, mouse and rat | Milteyi | 130-107-677 |
| Seahorse XFp Cell Mito Stress Test Kit | Agilent | 103010-100 |
| **Equipment** | | |
| Osmotic pumps | ALZET | Model 1007D |
| **Experimental Models** | | |
| Tg(Tert-GFP)22Brlt | David Breault | N/A |
| *mTert*-rtTA | David Breault | N/A |

| | | |
|---|---|---|
| B6.Cg-Tg(tet0-cre)1Jaw/J | The Jackson Laboratory | IMSR Cat# JAX:006234, RRID:IMSR_JAX:006234 |
| B6.129(Cg)-Gt(ROSA)26Sor$^{tm4(ACTB-tdTomato,-EGFP)Luo}$/J | The Jackson Laboratory | IMSR Cat# JAX:007676, RRID:IMSR_JAX:007676 |
| LSL-FLUC+/+ Rosa26 BL6J | The Jackson Laboratory | IMSR Cat# JAX:034320, RRID:IMSR_JAX:034320 |
| **Oligonucleotides** | | |
| For genotyping primer sequences see Table S1 | The Jackson Laboratory | N/A |
| qPCR *Cyclophilin* specific primer sequence, forward: CAAATGCTGGACCAAACACAA | N/A | N/A |
| qPCR *Cyclophilin* specific primer sequence, reverse: AAGACCACATGCTTGCCAT | N/A | N/A |
| qPCR *Gfap* specific primer sequence, forward: ACCAGCTTACGGCCAACAG | N/A | N/A |
| qPCR *Gfap* specific primer sequence, reverse: CCAGCGATTCAACCTTTCTCT | N/A | N/A |
| qPCR *Sox2* specific primer sequence, forward: ACCAGCTTACGGCCAACAG | N/A | N/A |

| | | |
|---|---|---|
| qPCR *Sox2* specific primer sequence, reverse: CCAGCGATTCAACCTTTC TCT | N/A | N/A |
| qPCR *Hes5* specific primer sequence, forward: GCACCAGCCCAACTCCAA | N/A | N/A |
| qPCR *Hes5* specific primer sequence, reverse: GGCGAAGGCTTTGCTG | N/A | N/A |
| qPCR *EGFR* specific primer sequence, forward: GCATCATGGGAGAGAAC AACA | N/A | N/A |
| qPCR *EGFR* specific primer sequence, reverse: CTGCCATTGAACGTACCC AGA | N/A | N/A |
| qPCR *DCX* specific primer sequence, forward: CATTTTGACGAACGAGAC AAAGC | N/A | N/A |
| qPCR *DCX* specific primer sequence, reverse: TGGAAGTCCATTCATCCG TGA | N/A | N/A |
| qPCR *NeuroD1* specific primer sequence, forward: ATGACCAAATCATACAGC GAGAG | N/A | N/A |

| | | |
|---|---|---|
| qPCR *NeuroD1* specific primer sequence, reverse: TCTGCCTCGTGTTCCTCGT | N/A | N/A |
| **Software and Algorithms** | | |
| Prism 9 | GraphPad | https://www.graphpad.com/scientific-software/prism/ |
| Zen 3.0 software | Zeiss Microscope | https://www.zeiss.com/microscopy/int/products/microscope-software/zen.html |
| NIS Elements | Nikon | https://www.microscope.healthcare.nikon.com/products/software/nis-elements |
| LAS X | Leica | https://www.leica-microsystems.com/products/microscope-software/p/leica-las-x-ls/ |
